# Dynamics of alpha suppression and enhancement may be related to resource competition in cross-modal cortical regions

**DOI:** 10.1101/2021.11.29.470447

**Authors:** Grace M. Clements, Mate Gyurkovics, Kathy A. Low, Diane M. Beck, Monica Fabiani, Gabriele Gratton

**Affiliations:** Beckman Institute, University of Illinois at Urbana-Champaign, IL, 61801, USA; Psychology Department, University of Illinois at Urbana-Champaign, IL, 61820, USA

**Keywords:** EEG alpha power, EEG theta power, alpha suppression, alpha enhancement, multimodal processing, resource competition

## Abstract

In the face of multiple sensory streams, there may be competition for processing resources in multimodal cortical area devoted to establishing representations. In such cases, alpha oscillations may serve to maintain the relevant representations and protect them from interference, whereas theta oscillations may facilitate their updating when needed. It can be hypothesized that these oscillations would differ in response to an auditory stimulus when the eyes are open or closed, as intermodal resource competition may be more prominent in the former than in the latter case. Across two studies we investigated the role of alpha and theta power in multimodal competition using an auditory task with the eyes open and closed, respectively enabling and disabling visual processing in parallel with the incoming auditory stream. In a passive listening task (Study 1a), we found alpha suppression following a pip tone with both eyes open and closed, but subsequent alpha enhancement only with closed eyes. We replicated this eyes-closed alpha enhancement in an independent sample (Study 1b). In an active auditory oddball task (Study 2), we again observed the eyes open/eyes closed alpha pattern found in Study 1 and also demonstrated that the more attentionally demanding oddball trials elicit the largest oscillatory effects. Theta power did not interact with eye status in either study. We propose a hypothesis to account for the findings in which alpha may be endemic to multimodal cortical areas in addition to visual ones.

## 1. Introduction

We constantly encounter complex sensory information from multiple sensory streams and must process this information to navigate the world. Regardless of whether the sights and sounds we perceive are relevant to us, in most cases they are processed at least to some extent, and may engender competition for processing resources. Two seemingly contradictory lines of research have investigated the oscillatory brain activity associated with this processing. On the one hand, in task-based electroencephalographic (EEG) recordings, an increase in the power of *pre-stimulus* posterior alpha oscillations (8-12 Hz) has been linked to the dampening of irrelevant information (e.g., Mathewson et al., 2009; Cosmelli et al., 2011; Vissers et al., 2016; for a review see Mathewson et al., 2011). The re-direction of attention to an unpredicted incoming stimulus is instead associated with the suppression of alpha activity (Feng, Störmer, Martinez, McDonald, & Hillyard, 2017) to allow for new representations to form. On the other hand, higher *post-stimulus* alpha power has been linked to increased memory for the stimulus itself (e.g., Jensen et al., 2002). As proposed by Gratton (2018), it is possible to accommodate both of these findings by assuming that increased alpha power represents a mechanism that helps the maintenance of existing representations in the presence of competing processing streams. Alpha activity is instead suppressed when new incoming stimuli need to be processed. In other words, whether alpha is suppressed or enhanced, and whether this is beneficial or detrimental to performance, depends on fine-grained dynamics that interact with the timing of incoming stimuli and the required processing. In this article we present a series of studies in which we examine these fine-grained dynamics in the context of multimodal processing.

### 1.1 The Role of Multimodal Competition

A long tradition in cognitive psychology refers to the processing of multiple stimuli as involving a competition for resources (Houghton & Tipper, 1984; Kahneman, 1973; Lavie, 1995; van der Heijden, La Heij, Phaf, Buijs, & van Vliet, 1988; Wickens, 1980, 2008). Indeed, it has been shown that stimuli compete for representation in the same cortical region (Desimone & Duncan, 1995; Reynolds, Chelazzi, & Desimone, 1999) and it is suggested that competition may underlie all resource limits (Scalf & Beck, 2010; Scalf, Torralbo, Tapia, & Beck, 2013). When multiple sensory systems, such as the visual and auditory modalities, input unrelated information, this competition would presumably occur in multimodal cortical regions, where it may lead to the visual and auditory signals attempting to establish competing representations in the same areas (e.g., Low et al., 2009). When our eyes are open, this may happen at any moment in time, as both the visual and auditory modalities can potentially feed new information at all times. In contrast, with closed eyes the visual processing stream is likely to be interrupted or suppressed at an early peripheral level and therefore unlikely to compete with the auditory modality in multimodal regions. A similar peripheral suppression is not necessarily easy to obtain for the auditory stream, since humans lack similar methods for switching off auditory input. We can therefore hypothesize that the maintenance of existing representations, the re-direction of attention, and the establishment of new sensory representations may differ when auditory stimuli occur while our eyes are closed compared to when they are open.

### 1.2 The Role of Alpha

In task-based settings, it is well established that alpha suppression occurs immediately after the presentation of both task-relevant and task-irrelevant stimuli across a variety of paradigms (e.g. Yamagishi et al., 2005; Thut et al., 2006; Foxe and Snyder, 2011; Vissers et al., 2016; Xie et al., 2016; Feng et al., 2017). As noted above, a possible interpretation of these findings is that alpha suppression facilitates the allocation of attention to a new stimulus by interrupting the ongoing maintenance of some previous representation. Accordingly, early alpha suppression tends to support task performance, facilitating the allocation of attention to a new incoming stimulus (Feng et al., 2017; Thut et al., 2006). Although less extensively researched than alpha suppression, some studies report alpha enhancement in a later time window, to maintain recently presented task-relevant information, such as during a working memory retention period (Jensen et al., 2002; Xie et al., 2016) or in a longer (∼1000 ms) interstimulus interval after a cue (Banerjee, Snyder, Molholm, & Foxe, 2011; Foxe, Simpson, & Ahlfors, 1998). In other words, this later alpha enhancement may serve to protect the newly formed representations from interference and is related to improved performance in these studies. These findings are integrated in a model proposed by Gratton (2018) in which alpha is part of an active neural system supporting the processing, maintenance, and updating of representations. This model purports that a representation’s initial processing may be facilitated by alpha suppression, after which its maintenance is protected by alpha enhancement.

The processing of representations may be especially challenging when multiple competing stimulus streams are present. As described above, one such situation is when (at least) two sensory modalities provide unrelated information at the same time, such as vision and hearing. Here we investigated whether suppressing visual input by closing one’s eyes impacts the processing of auditory information, as indexed by alpha activity. Specifically, we considered the following hypotheses. First, with closed eyes it is less likely that multimodal areas would be occupied by current visual representations when an auditory stimulus is presented. Therefore, less alpha suppression should be needed in this condition than when the eyes are open. Nonetheless, some alpha suppression should still occur when a new sound is presented. Of note, this prediction regarding stimulus-related alpha suppression occurs on top of the well-known general alpha reduction when the eyes open after being closed (e.g., Berger, 1929; Adrian & Mathews, 1934; Polich, 1997; Barry et al., 2007; Clements et al., 2021). Additionally, we hypothesize that it will be easier to generate and maintain new representations of auditory stimuli with closed than with open eyes because there will be little competition between vision and hearing, and that should be reflected in alpha dynamics. Specifically, if alpha activity represents a process by which representations are supported in multimodal cortex, one would be led to expect that auditory stimuli may in fact generate alpha enhancement following the initial suppression. Again, this would be particularly evident with eyes closed, because the higher-order cortical regions involved in cross-modal processing should be fully devoted to process the auditory representations under these conditions. However, if alpha activity is instead solely related to the processing/gating of visual information, then this alpha rebound should not be observed.

### 1.3 The Role of Theta

Although the current study focuses on posterior alpha activity, it can be expected that frontocentral theta activity (4-8 Hz) may also be involved in stimulus processing, given that theta bursts are thought to facilitate the redirection of attention to a new stimulus as cognitive control processes are engaged (Gratton, 2018; Gratton, Cooper, Fabiani, Carter, & Karayanidis, 2017; Landau, Schreyer, Van Pelt, & Fries, 2015; Sauseng, Hoppe, Klimesch, Gerloff, & Hummel, 2007). In such instances, theta bursts at anterior electrode sites can be thought to manifest a mechanism that interrupts ongoing processing and resets attentional weights to facilitate the processing of new representations (Cavanagh & Frank, 2014; Gratton, 2018; Voloh & Womelsdorf, 2016). Indeed, increased theta power has been reported in cognitive control and attention tasks, reflecting instances in which the attentional system encounters a change or an unexpected outcome (Cavanagh & Frank, 2014; Cavanagh & Shackman, 2015; Cavanagh, Zambrano-Vazquez, & Allen, 2012; Michael X. Cohen, 2014). Although often discussed in relation to conflict monitoring (Cohen & Donner, 2013), after commission of error responses (Valadez & Simons, 2018), and during task switching (Cooper, Darriba, Karayanidis, & Barceló, 2016; Sauseng et al., 2006) phasic theta bursts can more generally be considered a mechanism used to redirect attention to a behaviorally relevant stimulus.

As such, one would expect theta bursts to occur in many – if not all – studies in which attention is engaged, and possibly interact with eye status. In most cases, researchers have studied theta during tasks requiring visual attention. If theta indexes primarily a visual attention control mechanism, there may not be much theta in auditory tasks. Alternatively, theta could be a general mechanism that engages regardless of modality, rather than a mechanism that only redirects attention between stimuli or tasks within a modality. Even in this case, one could hypothesize that in auditory tasks with closed eyes, less theta activity may be observed in response to auditory stimuli, because the visual system is already effectively disengaged and therefore less redirection is needed to engage with the auditory stimuli. As a result, with closed eyes, we can hypothesize that less theta power will be needed to facilitate the redirection of attention than with open eyes. Alternatively, a final possibility is that theta may operate as an all-or-none mechanism, whose purpose is to interrupt any ongoing oscillations (such as alpha) that help maintain current representations. In such cases theta activity would not interact with eye status because these ongoing oscillations are always present (albeit to varying degrees with eyes open vs. eyes closed).

### 1.4 The Current Study

To test these hypotheses, we conducted two experiments that include an eye status manipulation to assess if modulations of oscillatory activities occur similarly in both eye conditions. Studies 1a and 1b investigate whether alpha suppression and theta bursts occur when auditory stimuli are not task relevant but may capture attention, and whether they do so differentially, depending on whether the eyes are open or closed. Study 2 replicates Study 1 and also investigates whether and how directed attention influences the modulation of alpha and theta during an auditory oddball task.

## 2. STUDY 1

### 2.1 Study 1a

#### 2.1.1 Method

##### 2.1.1.1 Participants

Participants were recruited from the Urbana-Champaign area and had no history of psychological or neurological conditions. Eleven younger (*M*age = 22, *SD _age_* = 3, 55% female) and 12 older adults (*M_age_* = 72, *SD _age_* = 4, 50% female) comprised the sample. Older adults were included to assess a potential age effect in oscillatory engagement after stimulus processing. However, the results failed to show significant age-related effects (**Supp. Figures S1 and S2**), which were therefore ignored henceforth, focusing on a total sample of 23 adults. The study received approval from the Institutional Review board at the University of Illinois at Urbana-Champaign, and all participants signed informed consent.

##### 2.1.1.2 Procedures and stimuli

Participants completed six experimental blocks during one EEG recording session. Three blocks were resting-state: one with eyes open, one with eyes closed, and one with eyes open but wearing an eye-mask to block visual input. The resting state data were used for other purposes and are reported elsewhere (Clements et al., 2021; Gyurkovics, Clements, Low, Fabiani, & Gratton, 2021). The masked data were inconclusive^1^ and are not described further. The other three blocks included the same eye conditions (open, closed, masked) but in each block 25 tone pips were randomly presented to participants with a 5-10 second interstimulus interval jittered to avoid possible entrainment. These extended interstimulus intervals were used to ensure that the pips were unexpected and to allow participants to return to a baseline level of processing. During these blocks, no response was required from participants, who were instructed to sit quietly and simply “take in” or “enjoy” the pips. Each block was 2-3 minutes long, depending on the random selection of interstimulus intervals, and block types were counterbalanced across participants.

During the eyes-open blocks, participants fixated on a white fixation cross on a light gray background. The pips were a 500 Hz sinusoidal tone of 75 ms duration. Pips were presented binaurally at 75% of maximum volume from two speakers that were positioned symmetrically behind the CRT monitor and out of the participants’ sight.^2^

##### 2.1.1.3 EEG Recording and Preprocessing

The recording session took place in an electrically and acoustically shielded room. EEG and EOG were recorded continuously from 64 active electrodes mounted in an elastic cap (Acti-Cap) using a BrainAmp recording system (BrainVision Products GmbH). EEG was recorded from the scalp electrodes and the right mastoid, referenced to the left mastoid, with off-line re-referencing to the average of the left and right mastoids. Two electrodes placed above and below the left eye were used to compute a bipolar vertical EOG derivation to monitor blinks and vertical eye movements, whereas two electrodes placed ∼1 cm away from the outer canthi of each eye were used to compute a bipolar horizontal EOG derivation to monitor saccades. Impedance was kept below 10 kΩ. The EEG was filtered online using a 0.5 - 250 Hz bandpass and was sampled at 500 Hz.

Offline processing of EEG was performed using the EEGLAB Toolbox (version: 13.6.5, (Delorme & Makeig, 2004) and custom Matlab 2019b scripts (The MathWorks Inc., Natick, MA, USA). A 30-Hz low pass filter was applied. The pip blocks were epoched into 3000 ms segments centered around the pip (including 1500 ms of EEG recording before and 1500 ms after pip’s onset). Epochs with amplifier saturation were discarded (less than .01% of all trials). Ocular artifacts were corrected using the procedure described in Gratton et al. (1983), based on the bipolar EOG recordings. After eye movement correction, epochs with voltage fluctuations exceeding 200 *μ*V were excluded from further analysis to minimize the influence of any remaining artifactual activity. If more than 20% a participant’s epochs were marked for rejection, they were visually inspected to determine if one or two faulty electrodes were the cause. If so, their traces were replaced with the interpolated traces of the neighboring electrodes and reprocessed to regain the lost epochs.

Time frequency representations of the data were then derived using Morlet wavelet convolution with Matlab scripts modified from Cohen (2014b) . Epoched data were fast Fourier transformed and multiplied by the fast Fourier transform of Morlet wavelets of different frequencies. Morlet wavelets are complex sine waves tapered by a Gaussian curve. Thirty logarithmically spaced wavelets between 3 and 30 Hz were used. The number of cycles of the Gaussian taper ranged between 3-10 and logarithmically increased as a function of frequency in order to balance the tradeoff between temporal and frequency precision.

An inverse Fourier transform was applied to the product of the FFT’d wavelets and the FFT’d data and power values were computed by squaring the length of this complex vector at each time point. To reduce edge artifacts during convolution, each epoch was tripled in length by using reflections on either side of the original epoch, such that the original epoch was sandwiched between two reflected versions of itself. Following time-frequency derivation, the reflected epochs were trimmed back down to their original length of 3000 ms.

Power values were baseline corrected using condition-specific subtractive baselining. We have previously shown that, compared to divisive baselining, subtractive baselining minimizes the potential of Type I errors that might occur due to the effect of the aperiodic, 1/*f* component of power spectra (Clements et al., 2021; Clements et al., in press, Gyurkovics et al., 2021). Baseline activity differs for the eyes open and eyes closed conditions, especially in the aperiodic 1/*f* activity (as well as oscillatory activity), such that the eyes-closed condition has a greater 1/*f* offset than the eyes-open condition (**Supp. Figure S1**). This difference could induce spurious effects (Type I errors), particularly at low frequencies. Given our interest in the difference between these two conditions, a subtractive baseline would mitigate errors induced by having differential baseline activities. The power in the baseline period (-1250 to -500 ms, chosen to minimize the influence of edge effects) was thus subtracted from the total power in each epoch.

##### 2.1.1.4 Statistical Approach

We were interested in investigating whether there was a differential alpha and theta engagement after hearing a passive pip with open eyes compared to closed eyes. Before assessing the difference between eye conditions, we tested whether alpha and theta differed from their baseline power levels. We chose to analyze the time-frequency space over a frontocentral subset of electrodes (Fz, FCz, Cz, CPz FC1, FC2, C1, C2) where theta is typically observed and a posterior subset of electrodes where alpha is typically the largest (Pz, POz, Oz, PO3, PO4, PO7, PO8, O1, and O2). These locations were informed by assessing the scalp topographies of activity compared to baseline and matched those used in a previous publication (Clements, et al., 2021). We used a permutation testing-based approach to assess the difference between baseline and post-stimulus activity as well as the difference between the eyes closed and eyes open time-frequency “maps” (i.e., a time x frequency heat plot representing, for each frequency [row] the power at each time point [column] relative to the average baseline value). In order to reduce computation time, we temporally down-sampled the time-frequency decomposed data to 40 Hz, such that we included power estimates every 20 ms, instead of every millisecond.

##### 2.1.1.5 Simple Effects

To appropriately use permutation testing, the user must define what the data would look like under the null hypothesis. If there was no difference from baseline after the pip, then the pre-stimulus and post-stimulus activity would be similar, and the distribution of difference scores should be statistical unchanged when the sign of those differences are randomly permuted. To create such a situation, a null distribution of 10,000 possible across-subjects average maps was created. The sign of the difference map was changed for half of the subjects chosen at random before computing the average. For each of these permutated maps, the maximum and minimum values across the entire map were saved, thus generating a distribution of possible minima and maxima obtained under the null hypothesis. Both maxima and minima were saved because we were conducting a two-tailed test, to encompass both power suppression and enhancement after the pip.

We then compared the values at each pixel of the actual observed map (averaged across individuals) to the distributions of maxima and minima expected under the null hypothesis. Pixels greater than the 97.5^th^ percentile in the maximum pixel distribution and pixels smaller than the 2.5^th^ percentile in the minimum pixel distribution at a particular time and frequency were considered as showing significant power enhancement and suppression, respectively. Note that this procedure effectively protects from map-wise alpha errors at a *α =* .05 level, accounting for the multiple comparisons, although it is likely to be overly conservative for frequencies and time points with reduced variance in power.

This procedure was conducted separately at the posterior and frontocentral locations. This was deemed appropriate since it was anchored to specific hypotheses on the effects for alpha and theta. Significant regions on the average time-frequency heat maps are denoted by a contour line on the original data indicating pixels with corrected *p*-values < .05. All time-frequency heat maps presented in this article use this convention.

##### 2.1.1.6 Main Effect of Eye Condition

A similar procedure was applied to the analysis of eye condition. However, before the permutation procedure was applied, *difference* maps between eyes open and closed were computed for each subject. Then the same procedure described above was conducted, again separately for the two electrode sets.

#### 2.1.2 Results

As mentioned, older and younger participants in Study 1a did not differ significantly in time-frequency maps at posterior or frontocentral electrodes (**Supp. Figures S2 and S3**) and were thus combined for all subsequent analyses (*n* = 23). We first assessed the simple main effects at the posterior and frontocentral electrode sites, separately for eyes open and eyes closed. The resulting time-frequency maps indicate the effect of the pip on theta and alpha. Interestingly, at posterior sites there was significant alpha suppression following the pip when the eyes were open but not when they were closed (**Figure 1C**). Instead, with eyes closed, a brief period of small and not significant alpha suppression was quickly followed by a period of significant alpha enhancement. Both alpha suppression and enhancement had a posterior scalp distribution, consistent with previous work on alpha (**Figure 1B**).^3^ As expected, at frontocentral electrodes there was a pronounced, significant theta burst following the pip with both eyes open and closed (**Figure 1A**). Scalp topographies show that both conditions produced a mid-frontal distribution, as expected (**Figure 1B**). At posterior sites, this theta activity was smaller, but still significant with both eyes open and closed (**Figure 1C**).

**Figure 1:**
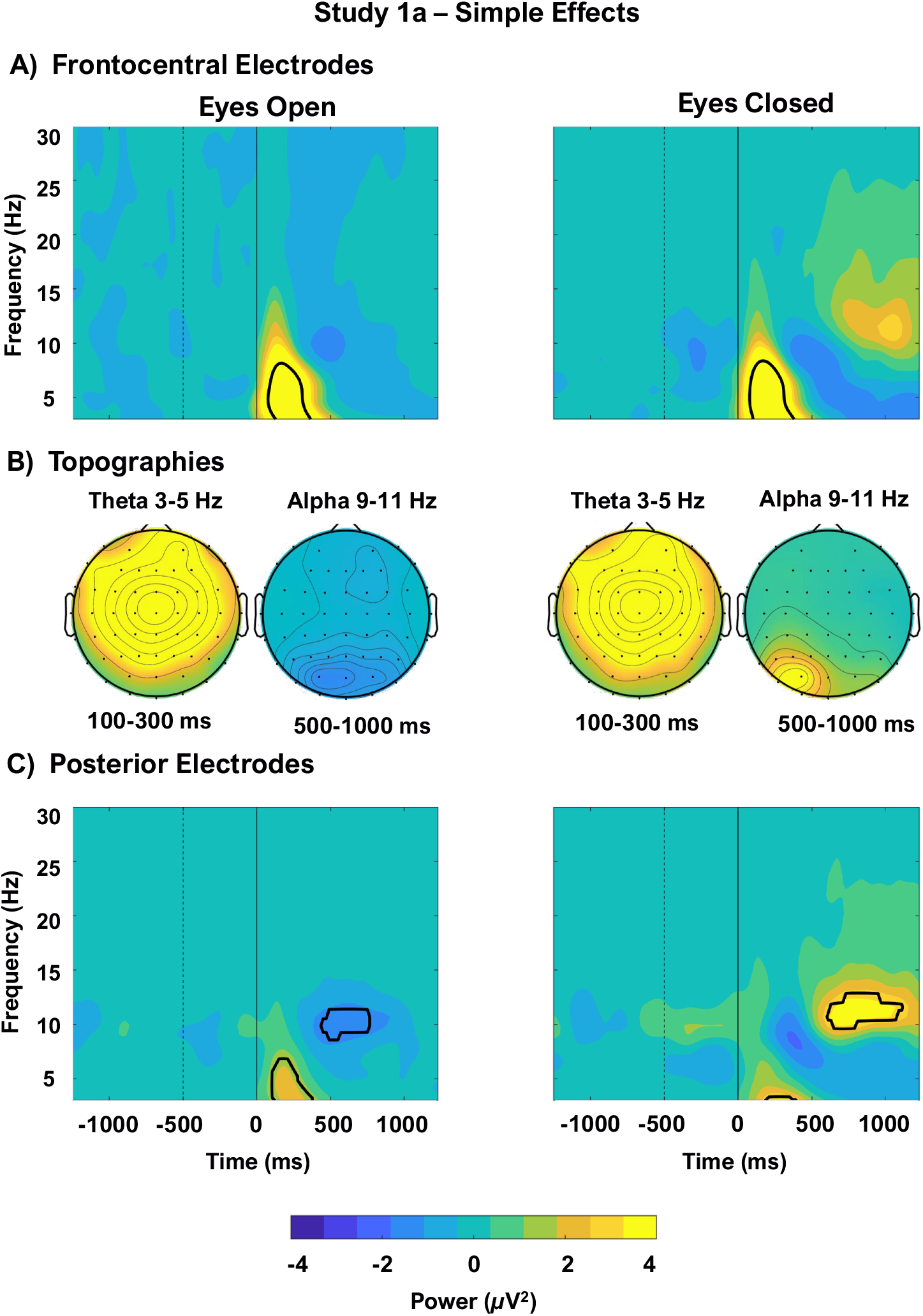
Simple effects (changes from baseline) for frontocentral (A) and posterior (C) electrodes following the pip with eyes open and closed for Study 1a (passive pips). The dotted vertical line indicates the end of the baseline period, the solid vertical line indicates stimulus onset. Black contours on the time-frequency maps outline significant pixels at *p* < .05, corrected for multiple comparisons. Note that significant theta activity occurred in all time-frequency maps, but it was more evident at frontocentral sites. At posterior electrodes (C) with eyes open, alpha suppression occurred; with eyes closed, both alpha suppression and enhancement occurred. (B) Scalp distributions of alpha and theta with eyes open and closed. All subplots are on the same scale.

These data indicate that, the pip elicited changes in both alpha and theta. Moreover, the pattern of activity appeared to be different for the two eye positions, at least at the posterior locations. We tested whether the open-eyes condition differed significantly from the closed-eyes condition at both frontocentral and posterior locations.

As mentioned earlier, at posterior electrodes, alpha suppression occurred after the pip when eyes were open (**Figure 2A**), but was much reduced when they were closed, being rapidly overtaken by a subsequent alpha enhancement (**Figure 2B**). Permutation testing confirmed this difference, showing greater alpha activity in the eyes closed condition that persisted from 500 to 1200 ms after pip onset (**Figure 2C**). Note that the significant region of the interaction observed in the heat maps presented in Figure 2C overlaps with both the late part of alpha suppression observed with open eyes and the period of alpha enhancement observed with closed eyes. This late alpha enhancement with closed eyes in response to pips has not been previously described. Therefore, to establish the replicability of this phenomenon, we conducted an exact replication of Study 1a with an independent sample of young adults, described next.

**Figure 2:**
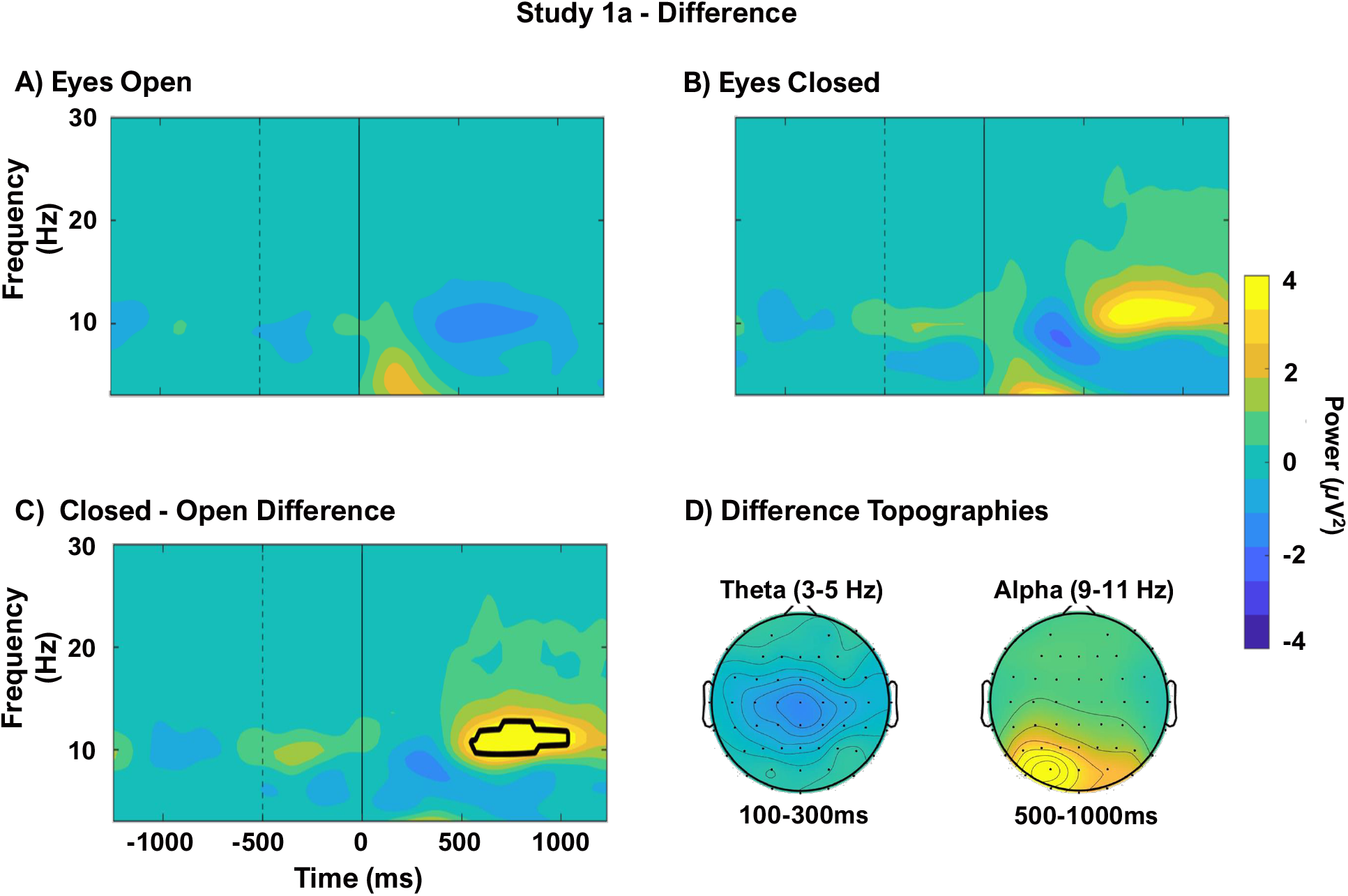
Comparison of the results obtained with eyes-open and closed from Study 1a at posterior electrodes. The dotted vertical line indicates the end of the baseline period, the solid vertical line indicates stimulus onset. Upper panels show the time-frequency responses after hearing a passive pip with eyes open (A) and eyes closed (B). Note that with eyes open (A) alpha suppression occurred and with eyes closed (B) alpha suppression was followed by alpha enhancement. (C) The difference between closed and open eyes was submitted to permutation testing and black contours outline pixels significant at *p* < .05, corrected for multiple comparisons. (D) Scalp distributions of the differences. All subplots are on the same scale.

The analysis at frontocentral locations did not show significant differences in the theta bursts elicited by the pips presented with eyes open or eyes closed (**Supp. Figure S4**). In combination with the simple effects, these data support the idea that theta bursts reflect a general process that does not vary with eye status.

### 2.2 Study 1b

#### 2.2.1 Method

Study 1b was a direct replication of Study 1a, with two exceptions. Because the results for the masked condition were inconclusive, this condition was not included in Study 1b. Similarly, as no differences had emerged between younger and older adults in Study 1a, for simplicity only younger adults were included in Study 1b. Participants were recruited from the Urbana-Champaign area and underwent the same EEG recording procedures as for Study 1a. Twenty-four younger adults comprised the sample, but one participant was excluded for not completing the task. The final sample consisted of 23 participants (*M _age_* = 22, *SD _age_* = 2.5, 61% female). EEG recording, preprocessing, and statistical approach were identical to Study 1a.

#### 2.2.2 Results

As in Study 1a, we replicated the finding of alpha suppression at posterior sites with eyes open and late alpha enhancement with eyes closed (**Figure 3C**). The alpha scalp distribution for both eye conditions was posterior and not lateralized, indicating that the left lateralized scalp distribution seen in Study 1a was likely a result of experimenter error with the speakers. There was also a significant theta effect after the pip for both eyes open and eyes closed at frontocentral locations (**Figure 3A**) and theta had a mid-frontal distribution in both conditions (**Figure 3B**). These results directly replicated the simple effects found in Study 1a.

**Figure 3:**
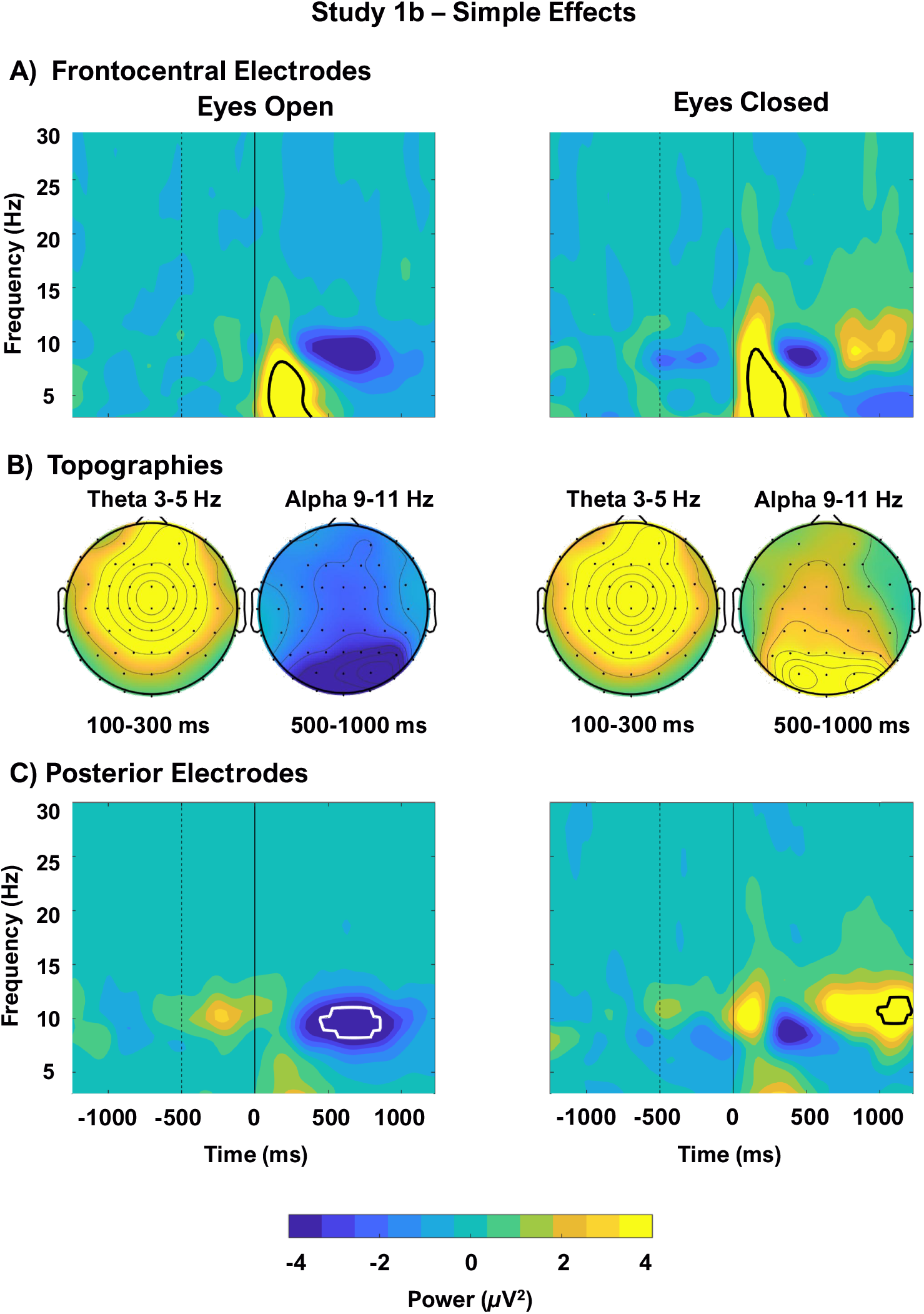
Simple effects (changes from baseline) for frontocentral (A) and posterior (C) electrodes following the pip with eyes open and closed for Study 1b (passive pip, replication). The dotted vertical line indicates the end of the baseline period, the solid vertical line indicates stimulus onset. Black contours on the time-frequency maps outline significant pixels at *p* < .05, corrected for multiple comparisons. Note that significant theta activity occurred at frontocentral electrodes (A) with eyes open and closed. At posterior electrodes (C) with eyes open, alpha suppression occurred and with eyes closed alpha suppression and enhancement occurred. (B) Scalp distributions of alpha and theta with eyes open and closed. All subplots are on the same scale.

As in Study 1a, we assessed the difference between eyes closed and eyes open using a permutation-based testing approach at posterior and frontocentral electrode sites. Again, the late alpha enhancement at posterior electrodes was greater with eyes closed relative to eyes open (**Figure 4 A-B**). The difference between open and closed eyes was significant between 500 – 1200 ms after the pip (**Figure 4C**). It encompasses both the alpha suppression with eyes open and the enhancement with eyes closed. There was no reliable difference in theta activity at frontocentral sites in this replication (**Supp. Figure S5**), providing further support that theta reflects a general process that is not modulated by eye status. The difference between closed and open eyes in Study 1b is nearly an identical replication of the effects in Study 1a, suggesting that alpha activity following a passive pip differs with eye status, but theta activity does not.

**Figure 4:**
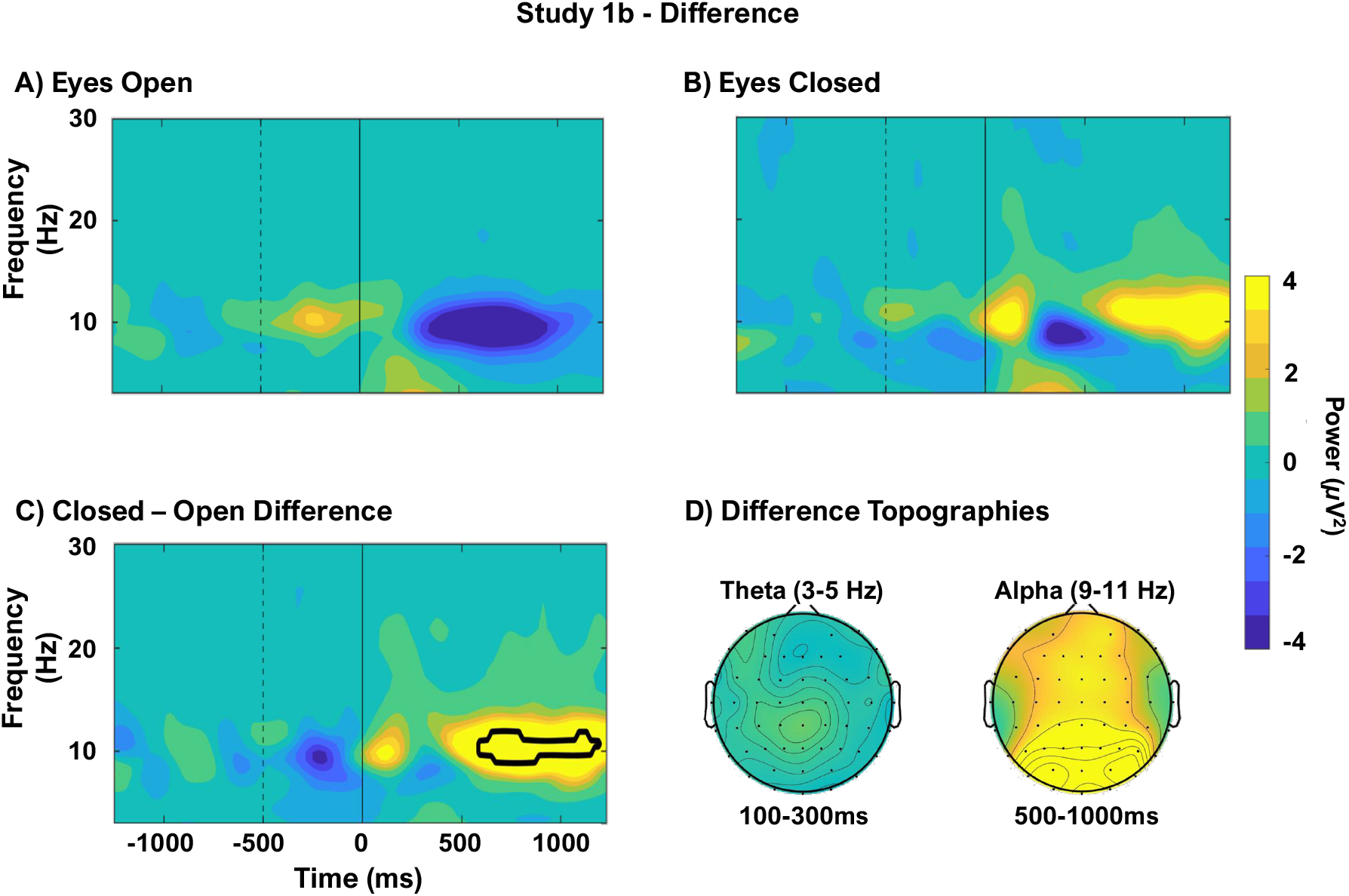
Comparison of the results obtained with eyes-open and closed from Study 1b at posterior electrodes. The dotted vertical line indicates the end of the baseline period, the solid vertical line indicates stimulus onset. Upper panels show the time-frequency responses after hearing a passive pip with the eyes open (A) and eyes closed (B). Note that with eyes open (A) alpha suppression occurred and with eyes closed (B) smaller alpha suppression was followed by alpha enhancement. (C) The difference between closed and open eyes was submitted to permutation testing and black contours outline pixels significant at *p* < .05, corrected for multiple comparisons. This difference is almost identical to that observed in Study 1a (Figure 2C). (D) Scalp distributions of the differences. All subplots are on the same scale.

### 2.3 Interim Discussion - Studies 1a and 1b

In the resting-state blocks with passive pips, the brain engages differently with open and closed eyes. Theta bursts immediately after the pip did not differ for eyes open and closed, indicating that redirection of attention to the sound occurs similarly for both eye states. Alpha suppression immediately followed the theta burst, and was more evident with eyes open than closed, suggesting that more attention may be needed to process the pip with eyes open than closed. These results suggest that the attention system is working harder when the eyes are open than when they are closed.

After the alpha suppression, we observed a late increase in alpha that was only evident with closed eyes. The replication study (1b) was conducted to provide more power and to further examine the alpha enhancement at long latency. Both studies 1a and 1b showed the same late alpha enhancement emerging around 500 ms in the closed-eyes condition. A possible interpretation for this late alpha enhancement is that closing the eyes frees up resources for processing the sound, presumably in multi-sensory cortical regions. These resources would instead be tied up with processing competing visual stimuli when the eyes are open. Note that this interpretation requires assuming that alpha activity is indeed the expression of the engagement of multi-sensory cortical regions. In turn, this requires considering that posterior alpha is not exclusively a reflection of visual processing but may reflect multi-sensory processing when information is present in multiple modalities.

If alpha suppression reflects attentional engagement, then we would expect it to increase when attention is explicitly required in an auditory task. To determine whether these effects observed during passive listening are affected, and perhaps even enhanced, by overt attention and active engagement in stimulus processing, we conducted a second study that includes an active auditory oddball task, described next.

## 3. STUDY 2

### 3.1 Method

#### 3.1.1 Participants, Procedure, and Stimuli

Participants in this study were the same as for Study 1b. After completing the conditions for Study 1b, participants completed an auditory 2-pip oddball task with eyes open and eyes closed. The two tone pips were a 500 Hz sine tone of 75 ms duration (identical to that used in Studies 1a and 1b) and a 450 Hz sine tone of 75 ms duration, and were randomly selected with an 80:20 (frequent:rare) probability and presented at 5-10 second interstimulus intervals. Participants were instructed to mentally count the rare tones (no movement or button press was required) and then report their total count to the experimenter at the end of each block. Given the long interstimulus intervals and to reduce fatigue, participants completed 12 blocks, each including 10 frequent pips and 2-3 rare pips. The blocks were approximately 3 min long, six with eyes open and six with eyes closed, for a total of 30 rare pips across the 12 blocks. Block order was counterbalanced based on eye status. For half of the participants, the rare pip was the same tone that they had heard in the passive pip blocks, for the other half, the frequent tone was the same as the passive pip. The same two tones were used for all participants.

#### 3.1.2 EEG Recording and Analysis

The EEG recording set-up and pip delivery system was the same as in Study 1a & b. The initial preprocessing and time-frequency processing steps were also identical to those in Study 1.

To assess sequential effects, the data were binned into the following trial-types: rare pips preceded by a frequent pip (“frequent-<u>rare</u>”, accounting for approximately 16% of the trials), frequent pips preceded by a rare pip (“rare-<u>frequent</u>”, also accounting for approximately 16% of the trials), and frequent pips preceded by a frequent pip (“frequent-<u>frequent</u>”, accounting for approximately 64% of the trials). Rare pips could also occasionally be followed by another rare pip (accounting for 4% of the trials), but these types of trials were not analyzed because these cases were extremely infrequent (and therefore yielded very noisy signals) and did not provide critical theoretical insights. The first pip in the block was also not used for the analyses because it was not yet in a sequence.

The frequent-<u>frequent</u> bin had more trials than the other two bins of interest (frequent-<u>rare</u>, rare-<u>frequent</u>). Therefore, to take into account the different number of trials for rare and frequent pips when assessing sequential effects, we randomly subsampled trials within each condition so that within each participant, the number of trials in each bin was equal to the minimum across the three trial-types.

In this study we derived and measured ERP waveforms in addition to time-frequency maps. The P300 obtained in an active oddball task is a well-documented index of task-relevant resource allocation and subjective stimulus probability (Donchin, 1981; Pritchard, 1981). There is an extensive and highly replicated literature linking the P300 elicited by oddball conditions to the allocation of attentional resources in counting tasks, and the reliability of these finding is sufficient to allow using this phenomenon as a method to assess the extent to which rare pips are processed ( Donchin & Isreal, 1980; Fabiani, Gratton, & Donchin, 1987; Squires, Donchin, & Squires, 1977). We calculated the mean P300 amplitude in the interval between 380-600 ms after the pip at the same posterior electrode set used in Study 1 (baselined to the average of the entire pre-stimulus period). Mean amplitudes for each participant, trial type and both eye conditions were calculated and submitted to a 2-factor repeated measures ANOVA in R (version 4.0.2; R Core Team, 2020). Normality was checked using the Shapiro-Wilk test and confirmed by examining the Q-Q plot. Follow-up paired *t*-tests of the simple effects were calculated, and *p*-values were Bonferroni corrected for multiple comparisons.

Similarly to Study 1, a permutation testing approach was used to analyze both the time-frequency simple effects between pre- and post-stimulus activity and the difference between eyes open and eyes closed for each of the three trial types at both posterior and frontocentral locations. The method of temporally down-sampling the data, generating the null map via 10,000 iterations and then pixel-based correction for multiple comparisons was identical, except that the three conditions were analyzed separately. The resultant time-frequency maps thus include pixels showing significant effects for each trial-type (frequent-<u>frequent</u>, frequent-<u>rare</u>, rare-<u>frequent</u>). Again, significant pixels with corrected *p*-values < .05 are denoted by a contour line on the time-frequency maps.

### 3.2 Results

#### 3.2.1 Behavior

Behavioral performance was assessed by calculating counting accuracy for the rare pips in the tone sequence. Accuracy was calculated for each block as 1 - abs(reported number of target/actual number of targets), and then averaged across blocks for each participant. Total accuracy for each participant (except for one who was missing accuracy data) was calculated across blocks and then averaged by eyes-open and eyes-closed conditions. Accuracy was similar with eyes open (*M* = .90, *SD* = .11) and closed (*M* = .89, *SD* = .10). Although neural engagement may vary between these conditions, closing the eyes did not affect simple counting performance. This may reflect the low level of task difficulty.

#### 3.2.2 ERPs

To assess sequential effects resulting from pip order, we measured the P300 at the posterior electrodes (**Figure 5**). Trial binning allowed us to examine whether simple changes in pip type across adjacent trials (occurring on both frequent-<u>rare</u> and rare-<u>frequent</u> trials, but not on frequent-<u>frequent</u> trials) or being a rare target (i.e., the difference between frequent-<u>rare</u> and rare-<u>frequent</u> trials) are relevant to the processing system. A 2 (eyes) × 3 (trial-type) ANOVA revealed a main effect of trial-type, *F*(2, 44) = 53.32, *p* < .001, generalized η^2^ = .362, but no effect of or interaction with eye status (*p* = .301, *p* = .907). Pairwise comparisons show that the P300 to frequent-<u>rare</u> pips was greater than rare-<u>frequent</u> pips *t*(45) = 8.68, *p* < .001, the P300 to rare-<u>frequent</u> pips was greater than frequent-<u>frequent</u> pips, *t*(45) = 3.59, *p* = .002, and that P300 to frequent-<u>rare</u> pips was also greater than frequent-<u>frequent</u> pips, *t*(45) = 10.5, *p* < .001 (Bonferroni adjusted *p*-values reported). These data indicate that when a change from frequent pip to rare pip occurs, which is the most relevant task change, more attentional resources are allocated to process the change than are required to process a change back from rare to frequent and even fewer resources are engaged when task-irrelevant frequent pips are presented consecutively.

**Figure 5:**
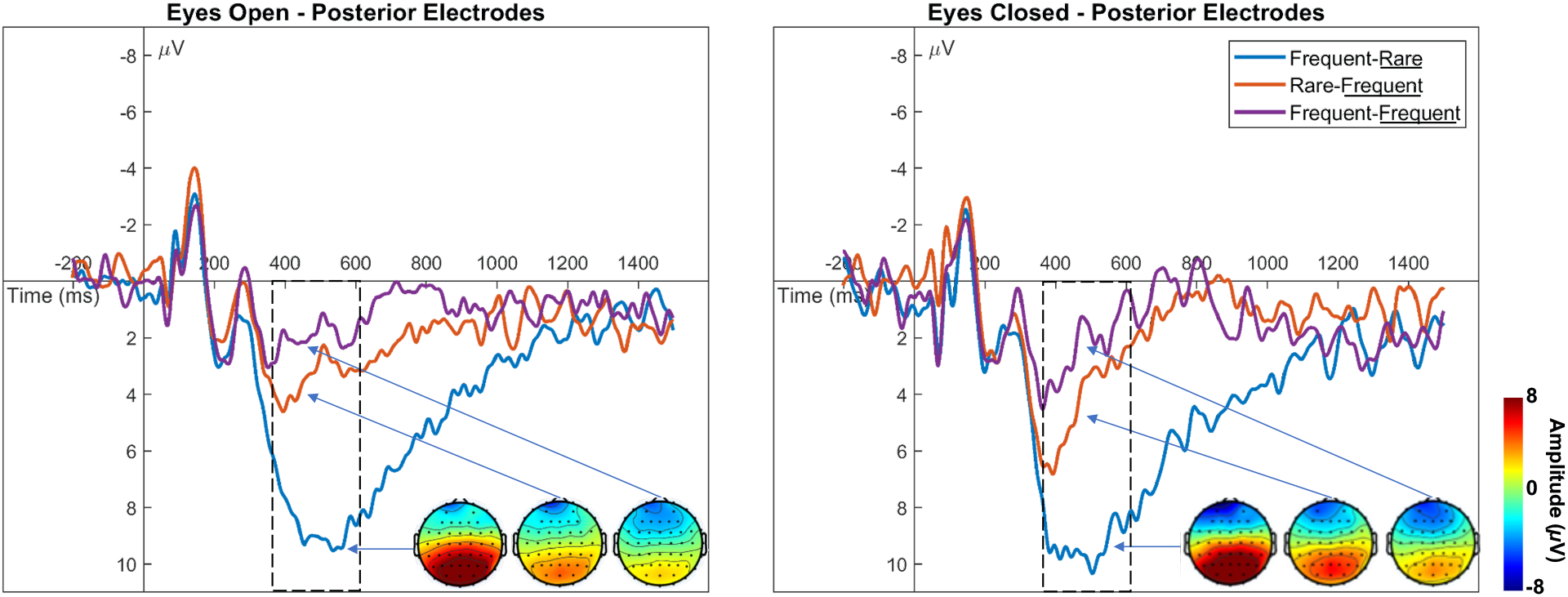
Grand-average ERP waveforms at posterior electrodes elicited by each of the oddball pip conditions (Study 2) with eyes open (A) and eyes closed (B). The dotted rectangle indicates the area under which P300 amplitude was measured. Scalp topographies for each trial-type are inset.

#### 3.2.3 Oscillatory Simple Effects

As in Study 1, simple effects were assessed based on eye status at posterior and frontocentral sites using a permutation-testing-based approach. Here, we assessed the effects of each condition in the oddball task. Once more, significant pixels with corrected *p*-values < .05 are denoted by contour lines on the time-frequency maps. With eyes open for all trial types, alpha suppression at posterior sites started at about 350 ms after the pip onset (**Figure 6A**). This effect appears strongest and longest in the frequent-<u>rare</u> condition, where it also had the broadest scalp topography, extending anteriorly. Pairwise comparisons between trial-types showed that frequent-<u>rare</u> pips had greater alpha enhancement than frequent-<u>frequent</u> pips but did not reveal significant differences between the other trial-types. (**Supp. Figure 6A**). These were calculated using the same permutation testing procedure described above and corrected for multiple comparisons with *p* < .05.

**Figure 6:**
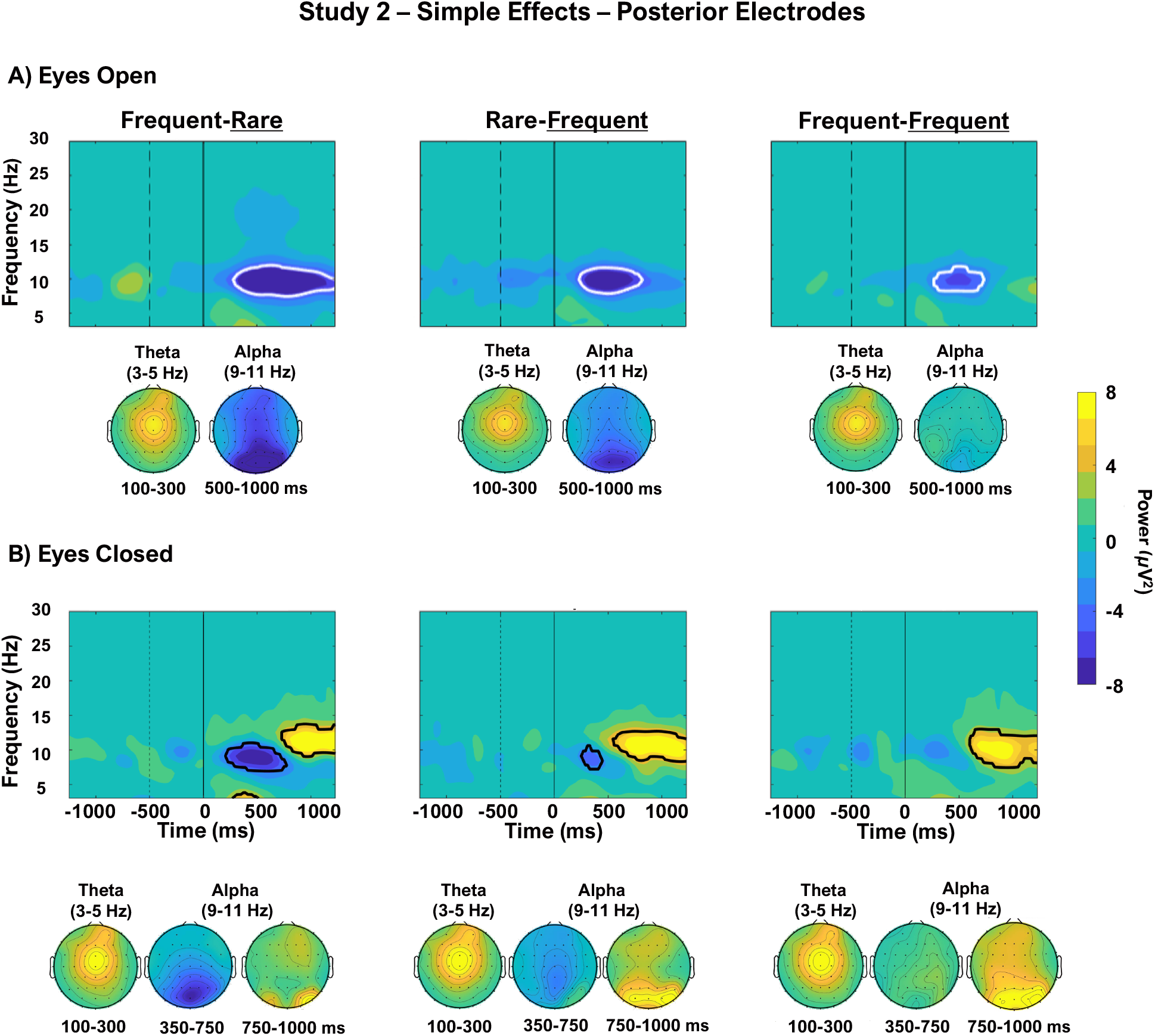
Simple effects (changes from baseline) for posterior electrodes during Study 2 (oddball). The dotted vertical line indicates the end of the baseline period, the solid vertical line indicates stimulus onset. Black contours on the time-frequency maps outline significant pixels at *p* < .05, corrected for multiple comparisons. (A) Eyes open time-frequency maps and scalp topographies ordered from the biggest effect (Frequent-<u>Rare</u> trials) to the smallest effect (Frequent-<u>Frequent</u> trials). Alpha scalp topographies are shown with two time-windows to illustrate the suppression (350-750 ms) and the enhancement (750-1000 ms). Note the robust alpha suppression with eyes open that diminishes in size across the trial-types. (B) Eyes closed time-frequency maps and scalp topographies. Note the alpha suppression followed by enhancement in the Frequent-Rare condition with eyes closed. All subplots are on the same scale.

With eyes closed, significant alpha suppression occurred in the frequent-<u>rare</u> and rare-<u>frequent</u> conditions (**Figure 6B**). In all three conditions, this was followed by a significant alpha enhancement. This pattern is consistent with the depth of suppression increasing with the level of engagement, as indicated by the P300 analysis. However, pairwise comparisons only detected differences between rare-<u>frequent</u> and frequent-<u>frequent</u> trials (**Supp. Figure 6B**). With the current sample size and the criterion used, we are unable to see the full gradation of effects across trial-types (**Supp. Figure 6C**). However, pairwise comparisons under both eye states indicate that trials in which a change occurred (frequent-<u>rare</u>; rare-<u>frequent</u>) have greater activity than no-change trials (frequent-<u>frequent</u>).

At frontocentral sites with eyes open for each of the three conditions, there was an initial theta burst (Figure 7A). This was followed by alpha suppression in the two change conditions, which appears longest in duration and largest in power on frequent-<u>rare</u> trials, slightly smaller, but still reliable on rare-<u>frequent</u> trials, and not significant on frequent-<u>frequent</u> trials. Pairwise comparisons support these assertions and indicate that alpha suppression was greater on both change trials compared to no-change trials (**Supp. Figure 7A**).

**Figure 7:**
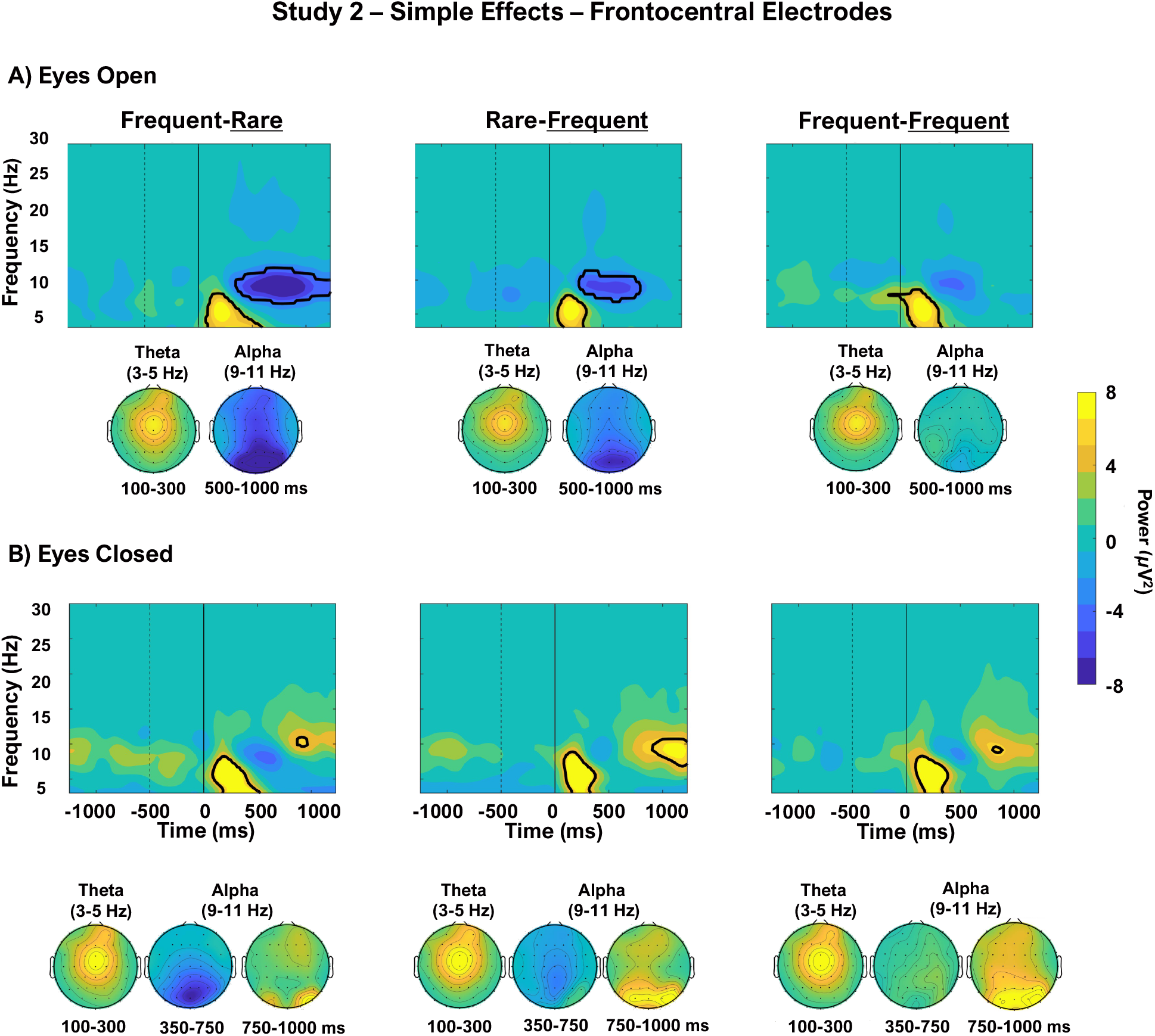
Simple effects (changes from baseline) for frontocentral electrodes during Study 2 (oddball). The dotted vertical line indicates the end of the baseline period, the solid vertical line indicates stimulus onset. Black contours on the time-frequency maps outline significant pixels at *p* < .05, corrected for multiple comparisons. (A) Eyes open time-frequency maps and scalp topographies ordered from the biggest effect (Frequent-<u>Rare</u> trials) to the smallest effect (Frequent-<u>Frequent</u> trials). Alpha scalp topographies are shown with two time-windows to illustrate the suppression (350-750 ms) and the enhancement (750-1000 ms). (B) Eyes closed time-frequency maps and scalp topographies. Note that significant theta activity occurs in all conditions. All subplots are on the same scale.

With eyes closed, theta bursts occurred in all three trial-types following the pips (**Figure 7B**). These bursts were followed by small, non-significant alpha suppression on frequent-<u>rare</u> trials, which was barely present in the other two conditions. All three trial-types showed reliable late of alpha enhancement. Pairwise comparisons again detected differences between change and no-change trials (Supp. Fig 7B). Although the effects appear to be in decreasing order in **Figure 7**, the current study was not powered to detect the full gradation.

Taken together, these data show that alpha suppression occurred at frontal sites with open eyes, but not with closed eyes, and at posterior sites with both eyes open and closed. Alpha enhancement was most evident at posterior sites with eyes closed, occurring in all three trial types.

Theta occurred at frontocentral sites with both eyes open and closed, but not posteriorly. As in Study 1, differential oscillatory engagement followed a pip, but in this case, this engagement resulted from processing task-*relevant* information. The differential engagement was distinct between change trials (frequent-<u>rare</u>; rare-<u>frequent</u>) and no-change trials (frequent-<u>frequent</u>).

#### 3.2.4 Oscillatory Differences in Eyes Open vs Closed

Time-frequency maps were subjected to permutation testing to assess the difference between eyes open and closed for each of the trial-types. As a reminder, we hypothesized that alpha suppression would occur following the pip with both eyes open and eyes closed, particularly at posterior sites. However, in Study 1, posterior alpha suppression was smaller with closed than open eyes. The oddball paradigm allowed us to determine if alpha suppression is smaller with closed eyes than open eyes in a task-relevant paradigm. We additionally hypothesized that theta may be reduced with closed compared to open eyes, particularly at frontocentral sites, indicating that with closed eyes less redirection of attention to the pip may be required than when the eyes are open. We will now discuss the effects at posterior and frontocentral locations in turn.

At posterior sites, alpha suppression was strongest with open compared to closed eyes (Figure 8A) and also for frequent-<u>rare</u> trials compared to frequent-<u>frequent</u> pips (**Supp. Figure 6** for pairwise comparisons). This suggests that after attention has been captured by the change to a rare pip, increased stimulus processing occurs. With closed eyes, some alpha suppression did occur at a similar latency, but it was much reduced compared to open eyes (**Figure 8B**).

**Figure 8:**
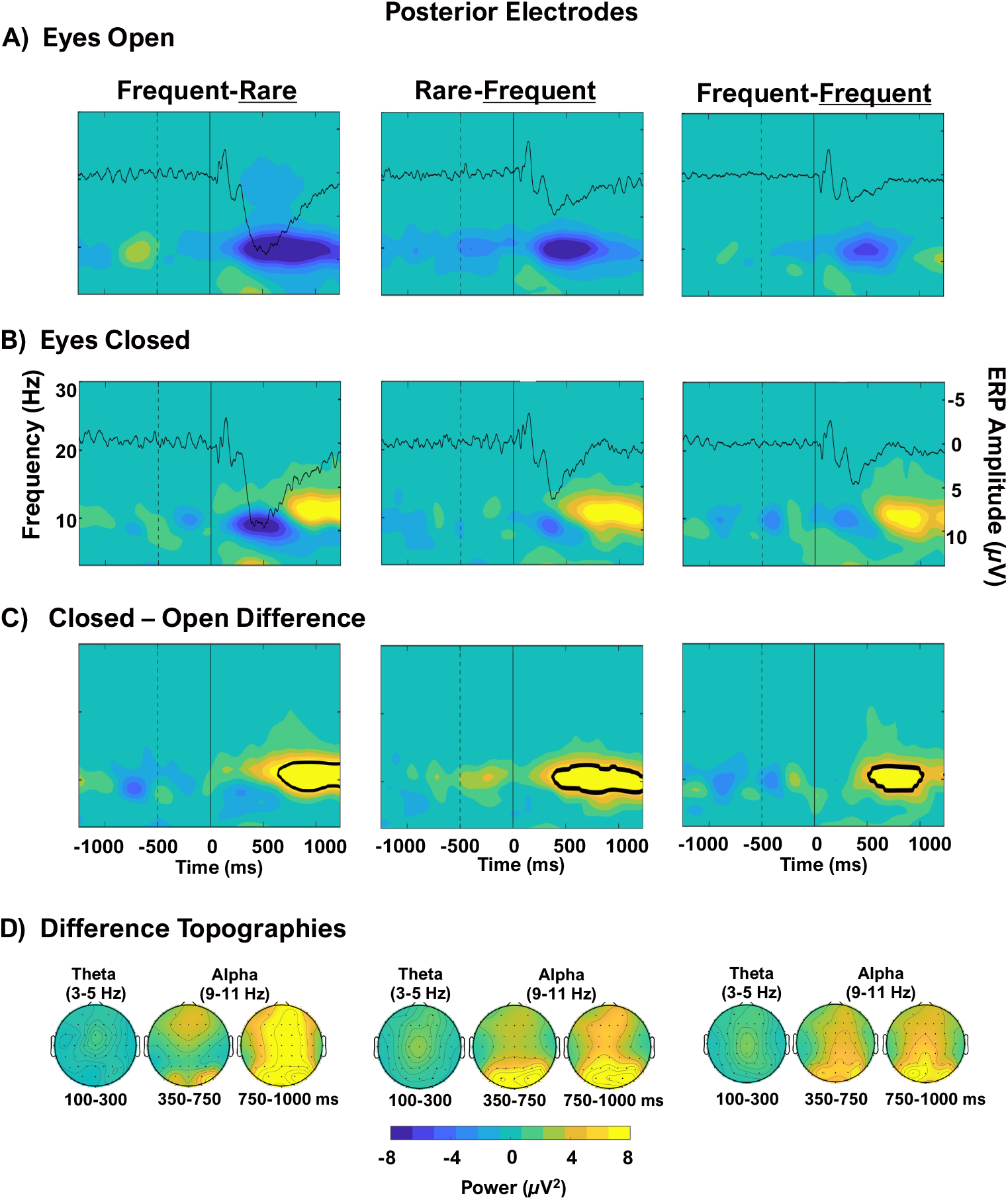
Comparison of the results obtained with eyes-open and closed from Study 2 (oddball) at posterior electrodes. The dotted vertical line indicates the end of the baseline period, the solid vertical line indicates stimulus onset. (A) Eyes open time-frequency maps ordered from the biggest effect (Frequent-<u>Rare</u> trials) to the smallest effect (Frequent-<u>Frequent</u> trials) with the average ERP at the same electrodes overlaid. (B) Eyes closed time-frequency maps with the average ERP overlaid. (C) The difference between closed and open eyes was submitted to permutation testing and black contours outline pixels significant at *p* < .05, corrected for multiple comparisons. (D) Scalp distributions of the differences. Alpha scalp topographies are shown with two time-windows to illustrate the suppression (350-750 ms) and the enhancement (750-1000 ms). All subplots are on the same scale.

Given the alpha enhancement found in Study 1 with eyes closed, Study 2 also sought to determine whether this effect could be modulated by attention using an oddball paradigm. As with the eyes closed passive pip blocks, alpha suppression was overtaken by alpha enhancement at posterior sites (**Figure 8B**). Indeed, there was a significant alpha difference between closed and open eyes beginning around 500 ms in all three trial-types (**Figures 8C**, with the significance contours denoting *p* < .05, corrected.) This difference encompasses both the alpha suppression occurring with open eyes (**Figure 8A**) and the alpha enhancement occurring with closed eyes (**Figure 8B**). This replicates the alpha enhancement found in Study 1 and extends it to conditions in which attention is actively engaged.

No theta differences were found at posterior sites between open and closed eyes, replicating the lack of an eye-status theta differences found in Study 1, suggesting that redirection of attention to the pip occurs similarly regardless of eye status (**Figure 8C**). Scalp topographies of the differences can be seen in **Figure 8D**.

A very similar pattern of effects was observed at frontocentral electrodes with one notable difference: the appearance of a theta burst in all conditions. While the alpha dynamics varied between open and closed eyes the same way as they did at posterior sites (**Figure 9C**), theta was not significantly different between open and closed eyes (**Figure 9 A-B**). Again, scalp topographies are shown in **Figure 9D**.

**Figure 9:**
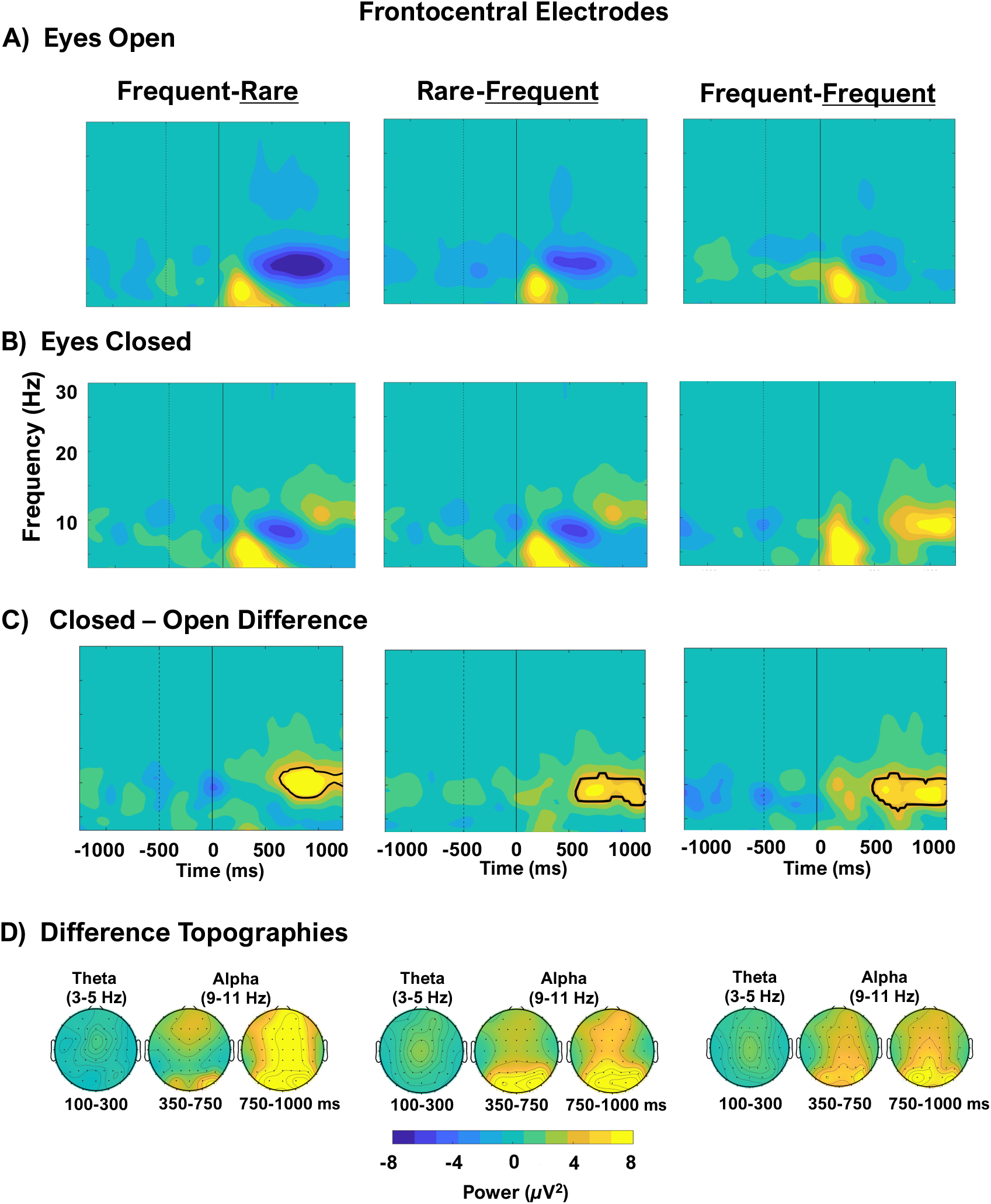
Comparison of the results obtained with eyes-open and closed from Study 2 (oddball) at frontocentral electrodes. The dotted vertical line indicates baseline onset, the solid vertical line indicates stimulus onset. (A) Eyes open time-frequency maps ordered from the biggest effect (Frequent-Rare trials) to the smallest effect (Frequent-Frequent trials) with the average ERP at the same electrodes overlaid. (B) Eyes closed time-frequency maps with the ERP overlaid. (C) The difference between closed and open eyes was submitted to permutation testing and black contours outline pixels significant at *p* < .05, corrected for multiple comparisons. (D) Scalp distributions of the differences. Alpha scalp topographies are shown with two time-windows to illustrate the suppression (350-750 ms) and the enhancement (750-1000 ms). All subplots are on the same scale.

## 4. General Discussion – Study 1 and 2

The two studies presented here indicate that auditory stimulus processing varies with eye status and with attentional engagement in response to pip frequency or by the amount of information provided by a stimulus. Alpha suppression occurred for both the passive (Study 1a, b) and the task-relevant tones (Study 2) but this suppression was more apparent in the eyes-open condition. The observation of greater alpha suppression with eyes open may show that more selection of the auditory stream is necessary with eyes open because the visual stream is competing with it. The hypothesis that alpha suppression is indexing the degree of selection or level of engagement is further corroborated by the larger alpha suppression following frequent-<u>rare</u> than frequent-<u>frequent</u> pips when the eyes are open. According to Gratton (2018), alpha suppression may be required to free up cortical regions (i.e., processing resources) from pre-existing sustained representations and make them available for subsequent processing. The observation that alpha suppression was elicited by auditory stimuli (pips) and was larger in the open eyes condition suggests that the cortical regions responsible for alpha might in fact include multimodal regions, which are sensitive to competition from other modalities beyond vision and may service at least auditory and visual stimuli.

Note that, in both studies, there was minimal alpha suppression when the eyes were closed. Instead, a later and broader alpha enhancement occurred in this condition. Again, this enhancement is suggestive of the idea that the alpha mechanism may be associated with multimodal cortical regions, in which competition may exist between processing streams from different modalities, and in which the ability to dedicate more cortical tissue to one modality (rather than sharing it across modalities) may result in enhanced processing (and the alpha enhancement we observed). Indeed, alpha enhancement to auditory stimuli has been observed between passive listening and task (Kolev et al., 1999) and with increasing task demands in an auditory oddball (Spencer & Polich, 1999). In other words, with the eyes closed, after a brief alpha suppression following the onset of a new pip to destabilize previous auditory representations, alpha was engaged again to protect the newly formed auditory representation. It is possible that the enhancement signal might be a “different alpha” with different generators from the suppression given their slight scalp topography differences. This should be further investigated in future work with different tasks. With open eyes, instead, more alpha suppression was needed to allow for the processing of the incoming stimulus, so that old auditory *and* visual representations can cease to compete with the new incoming stimulus. No alpha enhancement was observed in this condition. However, it is possible the extended alpha suppression with eyes open might have obscured any subsequent alpha enhancement, particularly if these responses may have different generators. In this sense, the oscillatory variability between eyes closed and eyes open during auditory processing may be interpreted as competition inherent to selective attention. With the eyes open, multiple sensory modalities are active and visual input may be competing with the auditory input to establish representations in the same, multimodal cortical region. Because of this, in order to process the pip, resources may be engaged to a greater extent with the eyes open than would be needed with the eyes closed.

The data are consistent with the proposal that theta bursts manifest the redirection of attention towards auditory stimuli (pips) even when they are passively listened to (Study 1a, replicated in Study 1b) and this occurs at similar levels irrespective of whether the eyes are open or closed. In fact, we show robust theta bursts at frontocentral sites with open and closed eyes to both passive pips (Study 1a, b) and during the oddball task (Study 2). Based on the alternative hypotheses presented in the introduction, this suggests that (a), theta activity occurs in both passive and attentionally demanding tasks as a way to redirect attention to whatever stimulus is presented, and (b) theta represents a more general mechanism that not only is engaged when switching between tasks/trials during visual paradigms (to begin a cascade of cognitive control processes, as shown in previous work, (e.g., Cavanagh & Frank, 2014; Gratton et al., 2017) but also is engaged even in simple, passive, auditory tasks and is independent of eye status.

In Study 2, as expected, we found that attention modulates the P300 amplitude, such that the most attentionally relevant pip sequence – a change from frequent to rare – resulted in the largest P300. This effect was graded such that the least attentionally relevant change (a frequent pip followed by another frequent pip) showed the smallest P300. The graded amplitudes indicate that resource allocation and context updating were occurring in response to the most attentionally relevant pips. However, P300 amplitude did not significantly differ with eye status. This has been reported before in a similar auditory oddball task (Spencer & Polich, 1999). Although the oscillatory effects suggest that the cross-modal nature of the study – the competition between vision and hearing occurring with eyes open – influences how the brain reacts to relevant sounds, whatever processes generate the oscillations may be distinct from those that generate the ERPs.

The extant literature provides several alternative interpretations of alpha enhancement, which only partially overlap with the one supported by the current studies, so we will discuss them in turn. Enhanced upper alpha power has been linked with better performance when a task requires tonic alertness or sustained attention, such as during monotonous breath-counting, auditory detection, or sustained attention response tasks (Braboszcz & Delorme, 2011; Dockree, Kelly, Foxe, Reilly, & Robertson, 2007; Sadaghiani & Kleinschmidt, 2016). The current study could be viewed as a sustained attention task, given the 5-10 second interstimulus intervals – even in the passive pip condition – so alpha may come online to facilitate processing the pip. Within this framework, it could be that alpha enhancement occurs as vigilance wanes. When the eyes are closed, participants may be more likely to drift off as attention dwindles. At pip onset, attention is refocused, and alpha engages to try to establish a representation of the tone, whether relevant or not. As a result, alpha is enhanced to a greater extent with closed eyes than open. If it is assumed that vigilance and attention are more likely to wane with eyes closed, then this interpretation could explain our findings. We will explore this hypothesis in future work as well as assess whether other stimuli elicit alpha enhancement.

Additionally, alpha enhancement has been related to motor response inhibition: perhaps with eyes closed there could be an impulse to open the eyes and orient to the pip (Mostofsky & Simmonds, 2008; Öhman, 1979). The enhancement would then be a result of inhibiting this impulse. However, given that alpha enhancement occurs late in the epoch (>500 ms) this explanation is not likely because an orienting reflex would occur shortly after the tone.

Based on the current studies, we propose that alpha oscillations are related to managing representations in *multimodal* cortical regions (or minimally, visual and auditory regions) or involved in a system that manages multimodal representations, and how and when alpha is engaged or suppressed depends on the dynamic requirements of the paradigm. Under this hypothesis alpha suppression interrupts the ongoing sensory stream to let new representations into multimodal regions for processing. Then, alpha enhancement occurs to form and maintain the new representation for processing. This is consistent with the idea of competition between processing streams (Kahneman, 1973; Treisman & Davies, 1973) and with the suggestion that alpha (and also beta) are generalized mechanisms used by the entire cortex (Gratton, 2018). In addition to the oft reported beta responses to movement (e.g., Little et al., 2019), beta enhancement has been observed in response to auditory stimuli in the absence of intended movement (Makeig, 1993; Fujioka & Ross, 2017; Fujioka et al., 2012) suggesting that its role may be similar to that of alpha within this framework. The alternative is that alpha oscillations are related to managing representations in visual cortical regions only and are therefore modality specific. In this case, the interaction with hearing emerges from another, connected, neural system. Future work will investigate these hypotheses to determine if alpha enhancement can be observed as a result of attention allocation to the visual or auditory modality during tasks with eyes open.

Some limitations should be pointed out. First, these studies used simple tasks in which behavioral performance, if any was required, was at ceiling. It is unclear, however, if these dynamic alpha effects would be similarly observed in more challenging tasks. However, the long ISIs that contributed to these tasks’ simplicity may be the reason we could observe the late alpha enhancement, because it came online 500 ms post-stimulus, which could be the beginning of the next trial in more fast-paced experiments. The current studies measured the impact of auditory processing with a visual manipulation, but ideally the converse of visual processing with an auditory manipulation would also be included. Some creativity is required to design an experiment that provides the same sensory experience with “ears closed” as with eyes closed. However, a visual task could be completed with varying levels of auditory stimuli using sound canceling headphones playing no audio vs. white noise. Future research should also investigate whether there are functional or behavioral consequences to the graded alpha enhancement following suppression.

In conclusion, across two studies, we showed that alpha activity varied dynamically in response to an auditory stimulus, changing with eye status and attention. Alpha suppression followed the typical pattern and occurred after both passive and relevant pips with both eyes open and closed, but it was greater when the eyes were open. With closed eyes, a later alpha enhancement occurred after alpha suppression in response to the passive pips. We replicated this effect in an independent sample (Study 1b) and then extended it in Study 2 using an attentional manipulation. Theta activity was elicited in both studies, primarily at frontocentral locations, but did not differ with eye status, suggesting that theta reflects a more general information processing mechanism. These results suggest that alpha activity may be endemic to, or may involve multimodal cortical areas as well as visual ones and future work should aim to further investigate this hypothesis.

## Supporting information

Supplemental Figs & Analysis

## Acknowledgments

This work was supported by NIA grant RF1AG062666 to G. Gratton and M. Fabiani. We acknowledge Daniel Bowie and Samantha Rubenstein for help with data collection.

## Footnotes

1 We could not definitively determine whether participants indeed had their eyes open during the recording. Several participants told the experimenter that they could not tell if their eyes were open or closed because the mask blocked out all light from the surroundings, as the recording occurred in a dimly lit room.

2 After assessing scalp topographies, it became clear that one of the speakers was not turned on for all sessions, resulting in a unilateral pip-presentation in some participants. This is one reason we chose to conduct a replication study (Study 1b).

3 As mentioned above, the alpha effects had a left lateralized scalp topography because for some participants one speaker used to present the pips was not turned on. This was addressed in the replication (Study 1b).

