## Supplemental Figs & Analysis for "Dynamics of alpha suppression and enhancement may be related to resource competition in cross-modal cortical regions"

Supplementary Materials

*Figures*

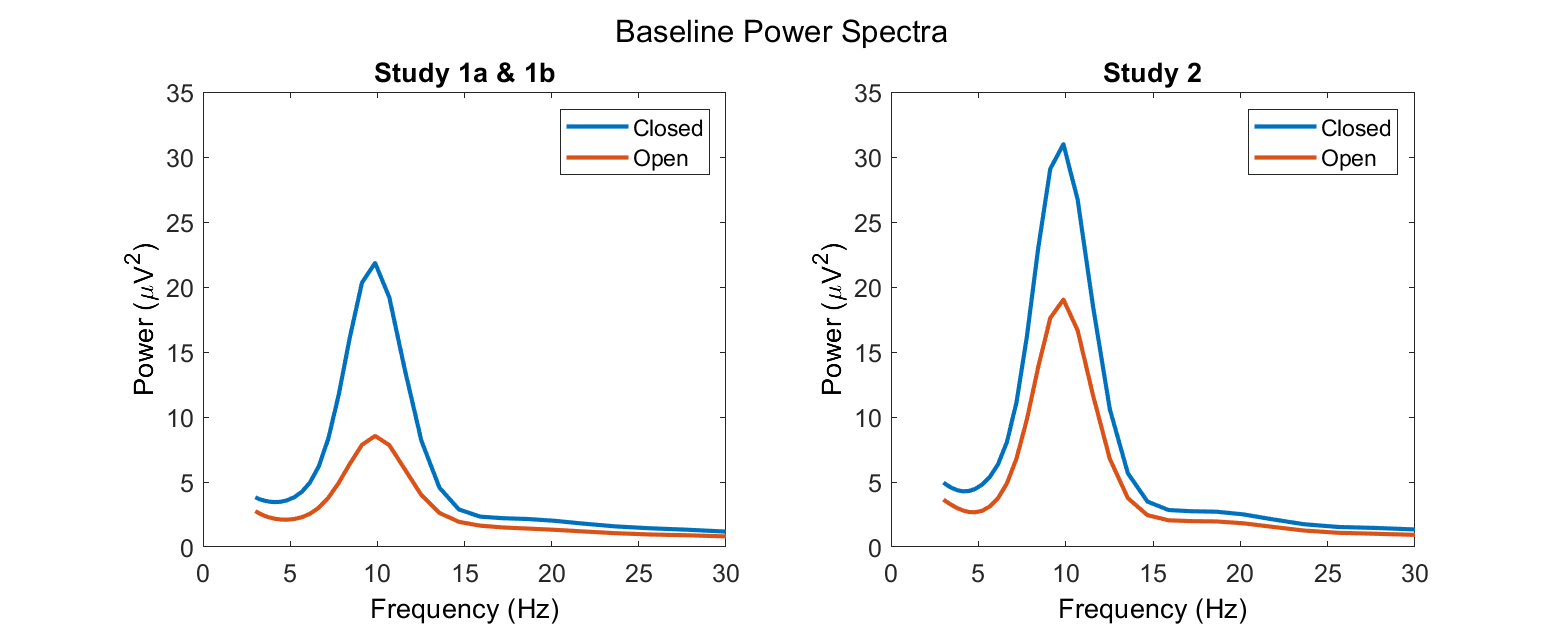

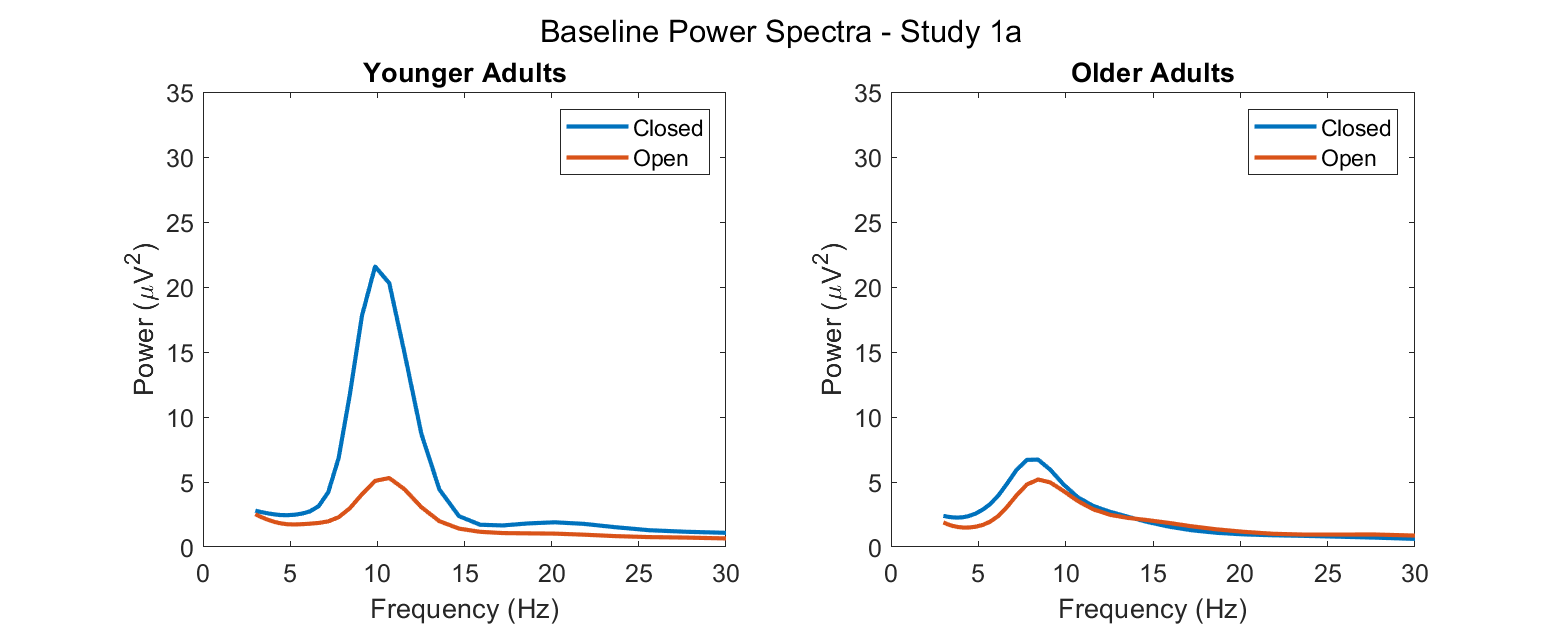

**Figure S1**: Top row: Power spectra of the baseline period (-1250 to -500 ms) for eyes open and eyes closed. These spectra are averaged over the posterior electrodes used in this study. In addition to a clear difference in alpha power, the offset of the 1/*f*-like component also differs between closed and open eyes in both Study 1a and 1b as well as Study 2. For this reason, a subtractive baselining procedure was used following wavelet convolution.
Bottom row: Power spectra of the baseline period for eyes open and eyes closed for the younger and older adults in Study 1a.

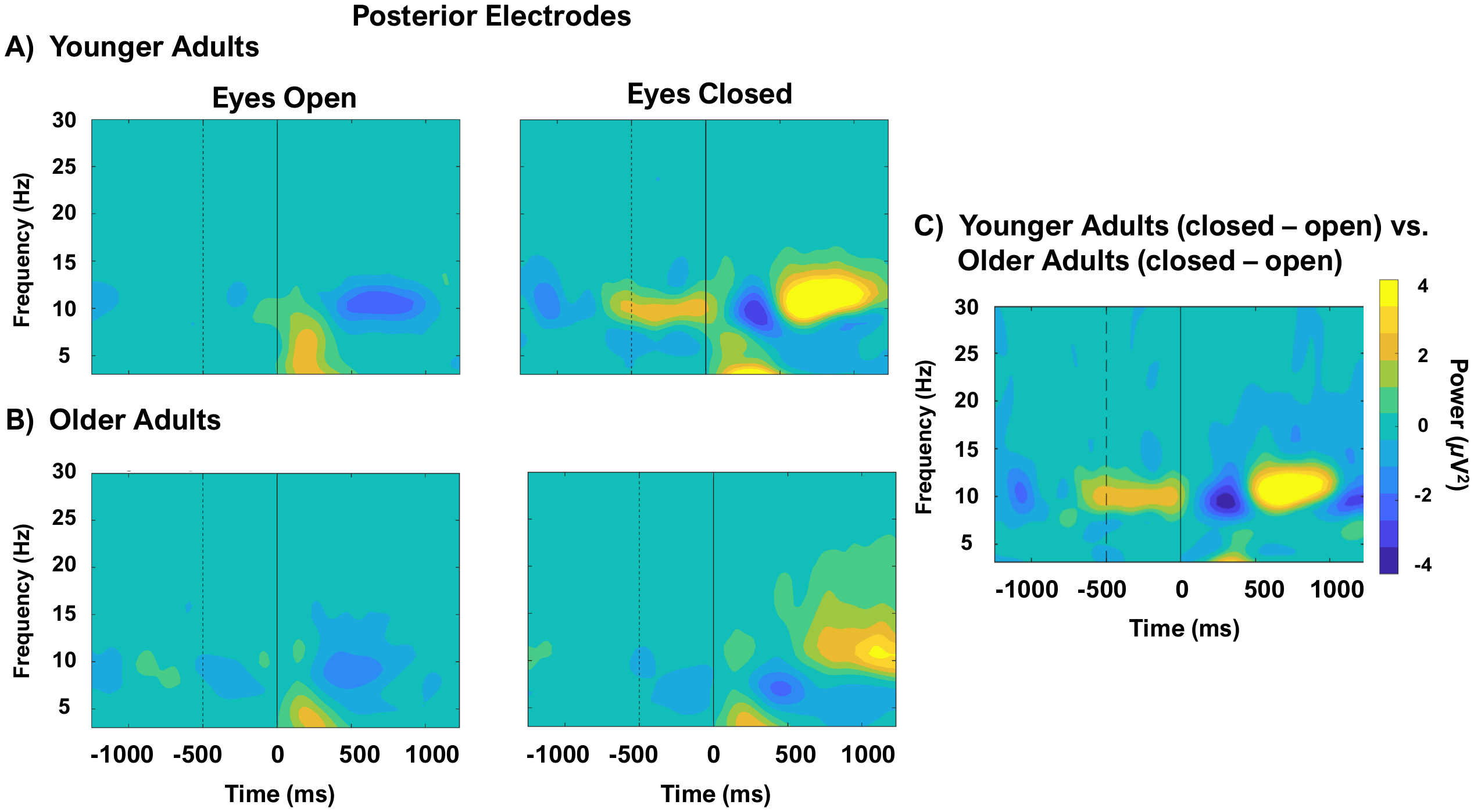

**Figure S2**: Time-frequency maps for the younger (A, n = 11) and older adults (B, n = 12) in Study 1a at posterior electrodes. The dotted vertical line indicates baseline ending, the solid vertical line indicates stimulus onset. With eyes open there is an initial weak theta burst followed by alpha suppression. With eyes closed there is also a weak theta burst followed by alpha suppression and then strong alpha enhancement. C) A permutation test of the difference between the younger and older adults’ closed-minus-open differences (age by eye status interaction) does not reveal any significant differences (*p* > .05, corrected for multiple comparisons).

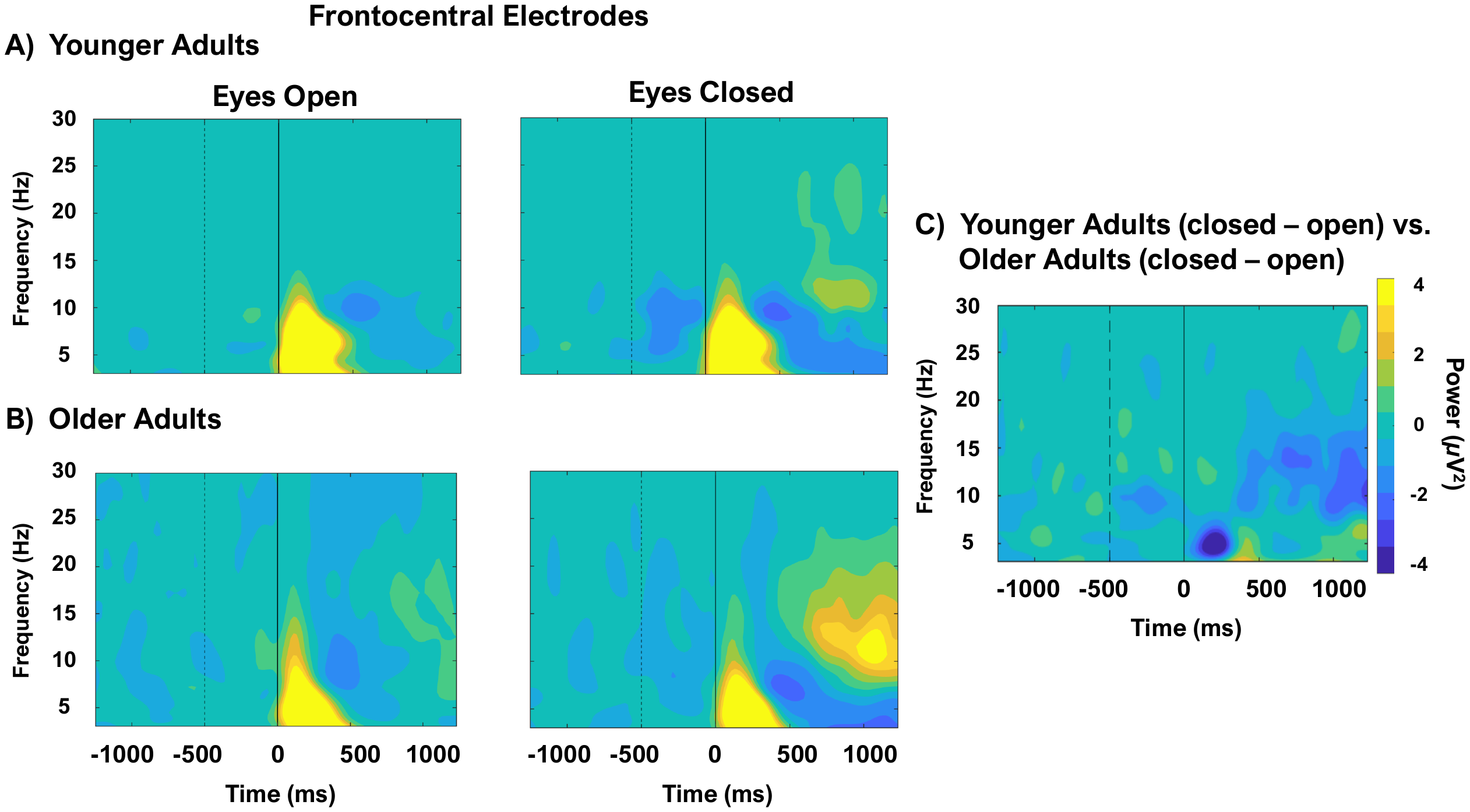

**Figure S3**: Time-frequency maps for the younger (A, n = 11) and older adults (B, n = 12) in Study 1a at frontocentral electrodes. The dotted vertical line indicates baseline ending, the solid vertical line indicates stimulus onset. With eyes open there is an initial strong theta burst followed by a smaller alpha suppression and with eyes closed there is also a strong theta burst followed by smaller alpha suppression and then alpha enhancement. C) A permutation test of the difference between the younger and older adults’ closed-minus-open differences (age by eye status interaction) does not reveal any significant differences (*p* > .05, corrected for multiple comparisons).

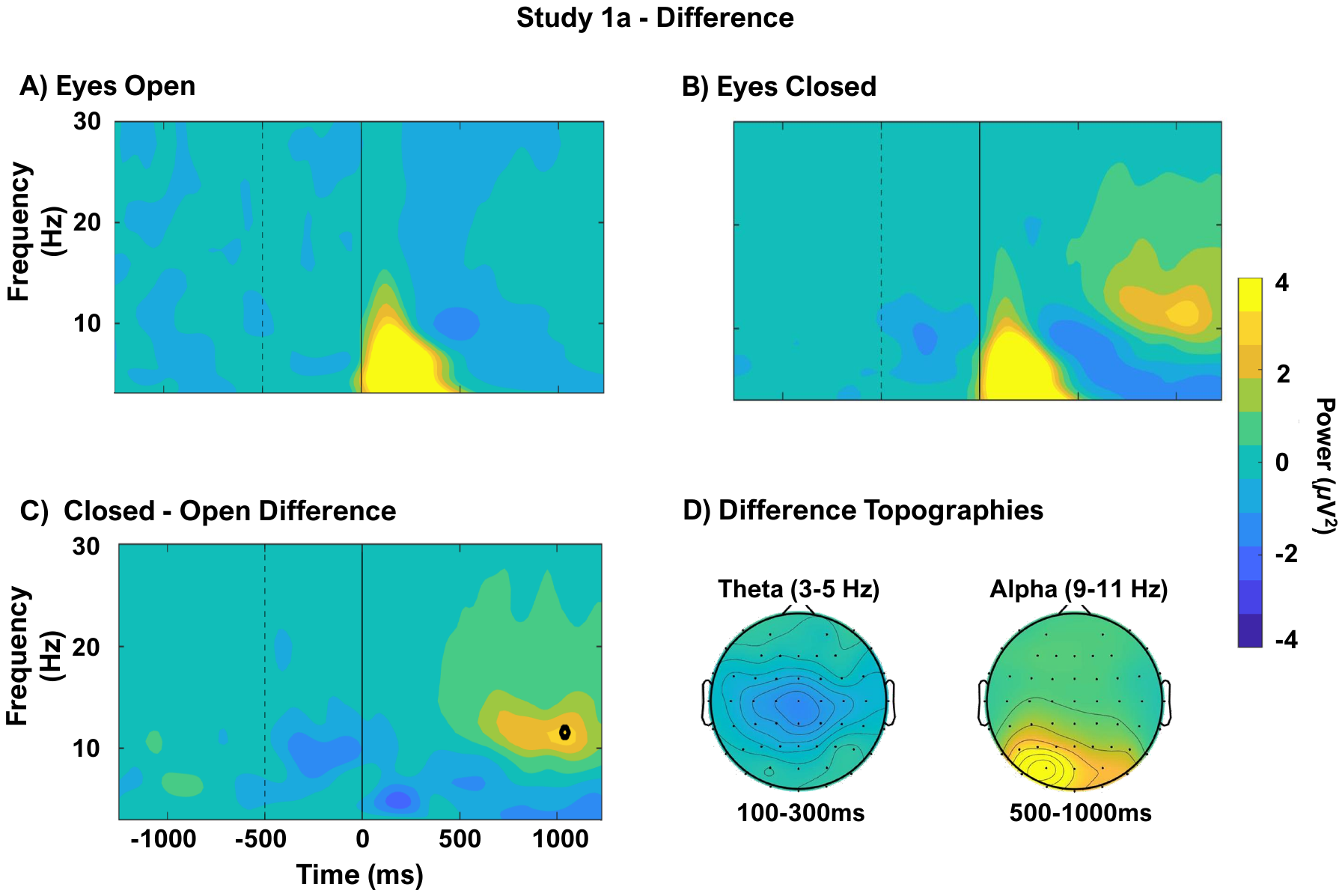

**Figure S4**: Test of the difference between eyes open and eyes closed at frontocentral electrodes in Study 1a, irrespective of age. The dotted vertical line indicates baseline ending, the solid vertical line indicates stimulus onset. There is no significant theta difference between these two conditions and only a few pixels in the alpha band pass the correction for multiple comparisons. Given our assessment of alpha at posterior locations, these pixels will not be considered further.

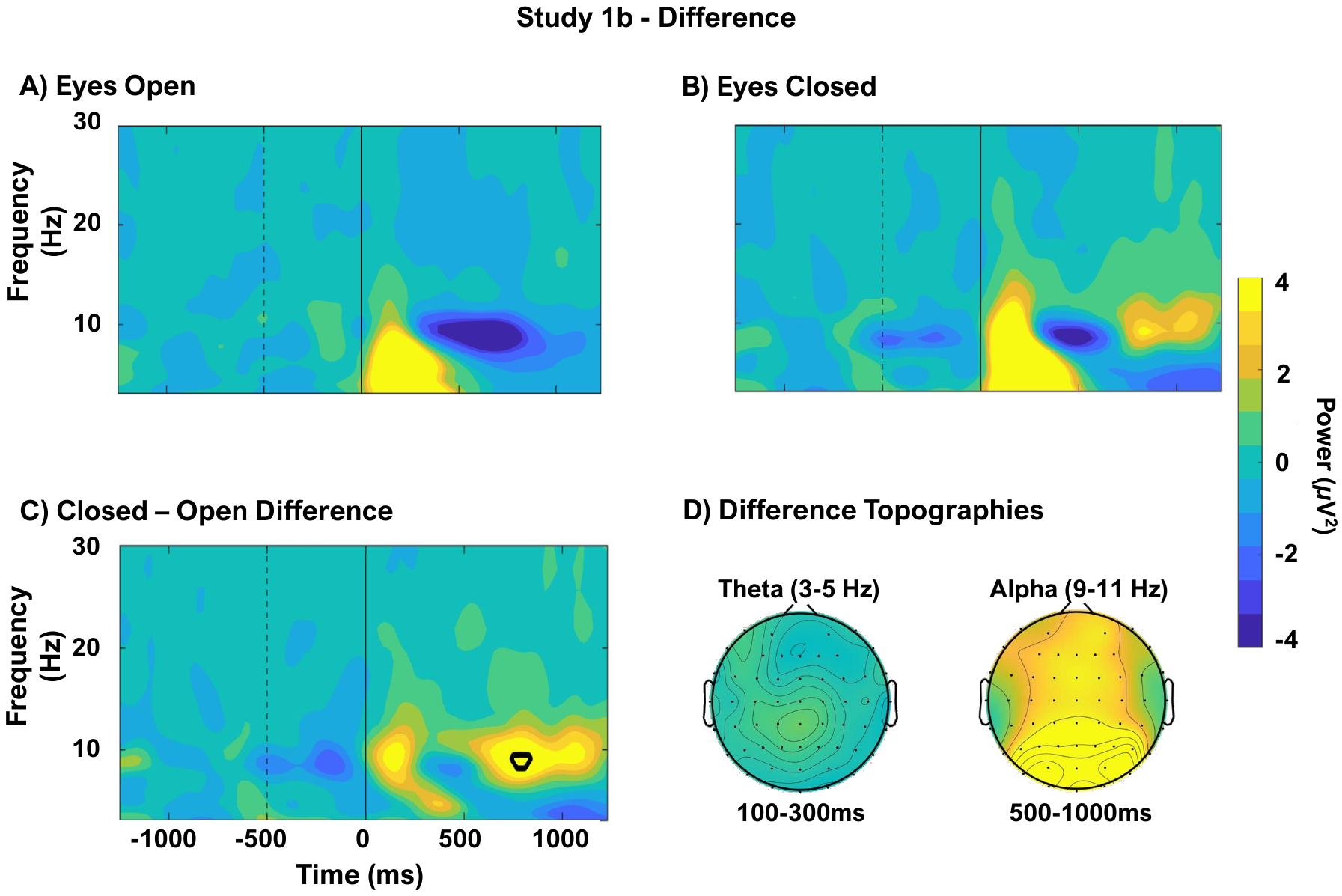

**Figure S5**: Test of the difference between eyes open and eyes closed at frontocentral electrodes in Study 1b. The dotted vertical line indicates baseline ending, the solid vertical line indicates stimulus onset. There is no significant theta difference between these two conditions and only a few pixels in the alpha band passed the correction for multiple comparisons. Given our assessment of alpha at posterior locations, these pixels will not be considered further.

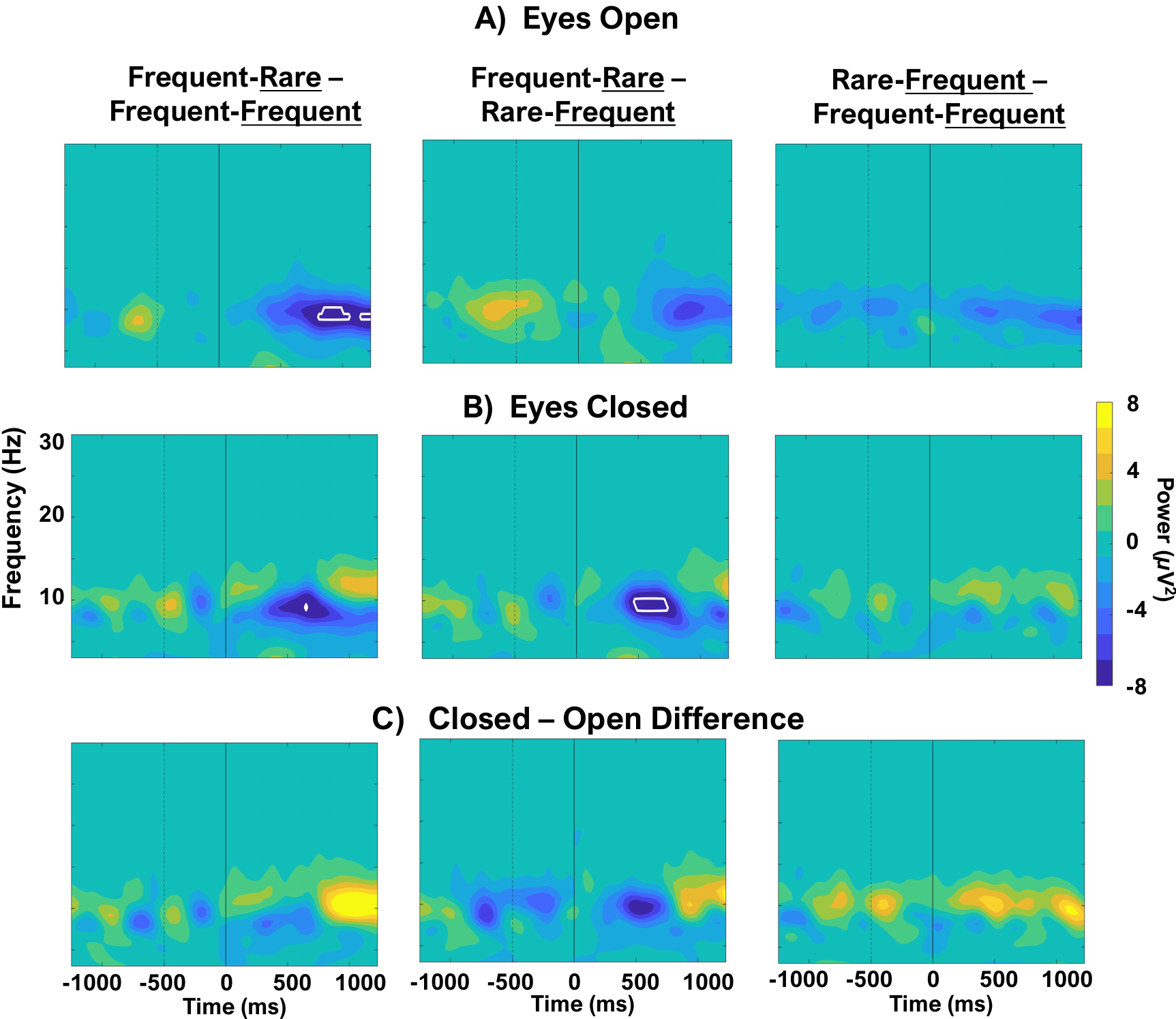

**Figure S6**: Pairwise comparisons across the trial types in Study 2 at posterior electrodes using a permutation testing-based approach. The dotted vertical line indicates baseline ending, the solid vertical line indicates stimulus onset. Each panel shows the time-frequency difference map between trial-types and contours outline pixels significant at *p* < .05, corrected for multiple comparisons. The eyes open (A) and eyes closed (B) pairwise comparisons show some significant differences between trial types in both eye conditions. However, the eye status by trial-type interactions shown in C are not significant (see discussion below).
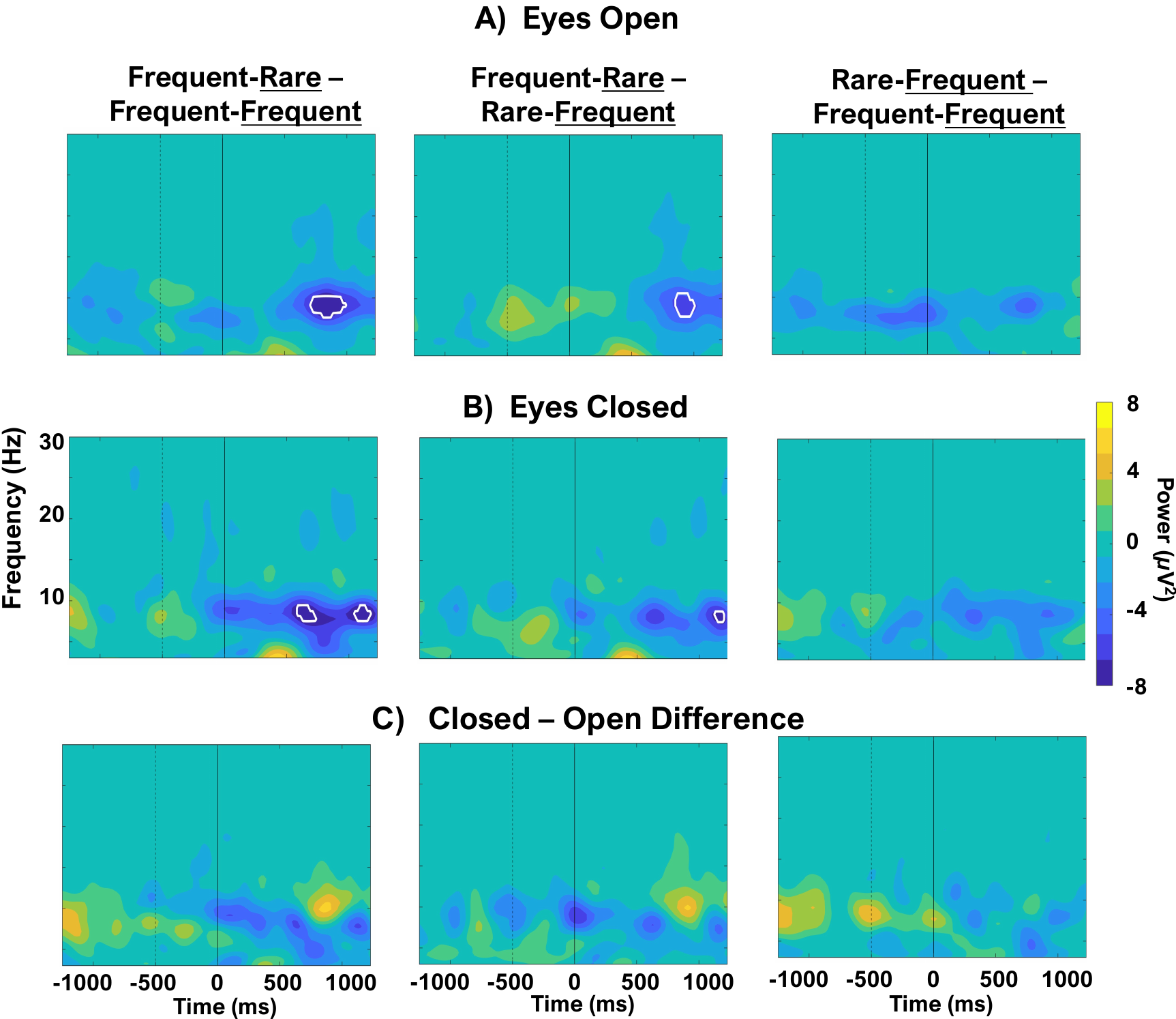

**Figure S7**: Pairwise comparisons across the trial types in Study 2 at frontocentral electrodes using a permutation testing-based approach. The dotted vertical line indicates baseline ending, the solid vertical line indicates stimulus onset. Each panel shows the time-frequency difference map between trial-types and contours outline pixels significant at *p* < .05, corrected for multiple comparisons. The eyes open (A) and eyes closed (B) pairwise comparisons show some significant differences between trial types in both eye conditions. However, the eye status by trial-type interactions shown in C are not significant (see discussion below).

*Ancillary Analyses*

Pairwise comparisons on the eyes closed-open difference for each trial-type was assessed at both electrode locations using a *p* < .05 criterion, corrected for multiple comparisons. This analysis assessed whether the eye status effect and the attention effect (if any) are additive or interactive. No differences were detected at posterior or frontocentral sites (**Supp. Fig. 6C & 7C**). However, at posterior sites an alpha trend was in the expected location (**Supp. Fig. 6C, leftmost panel**) – greater closed-open alpha band differences between frequent-rare and frequent-frequent trials beginning at 750 ms. This indicates that this is likely not random variability and there may be a true effect, but it is not substantial enough to be detected with the conservative criterion used (*p* < .05, corrected for multiple comparisons). Given these borderline results, we cannot conclude whether the eye status and the attention effects are interactive or additive. This should be tested in a larger sample.

To assess whether there was a significant interaction between the eye and trial type conditions, we employed a “double difference” permutation testing approach. We used the frequent-frequent trial-type as the baseline condition because little activity should be happening there. We then calculated the difference between frequent-rare and frequent-frequent separately for closed and open eyes, and then subtracted the difference between them:

$$Closed (FrequentRare - FrequentFrequent) - Open (FrequentRare-FrequentFrequent)$$

We did the same calculations for the rare-frequent trial-type (i.e., swapping out rare-frequent for frequent-rare in the equation above). These differences were subjected to permutation testing to determine whether the eye condition difference between frequent-rare and rare-frequent was significant. This is similar to testing the interaction term of a 2(eyes) × 3(trial-type) ANOVA. This was done at both the frontocentral and posterior sites. The same multiple comparisons corrections as in Study 1 were applied to the resultant time-frequency maps.

Using this approach, there were no significant differences in alpha or theta activity between the closed (frequent-rare) - open(frequent-frequent) effects found at frontocentral or posterior electrodes. We also did not find alpha or theta differences for the rare-frequent trials at either electrode location.

*References*

Davis, S. W., Dennis, N. A., Daselaar, S. M., Fleck, M. S. & Cabeza, R. (2008). Qué PASA? The Posterior–Anterior Shift in Aging. *Cerebral Cortex 18(5)*: 1201–1209. <https://doi.org/10.1093/cercor/bhm155>

Koleva, V., Yordanovaa, J. Y., Basar-Eroglu, C., & Basar, E. (2002). Age effects on visual EEG responses reveal distinct frontal alpha networks. Clinical Neurophysiology, 113(6), 901-910. <https://doi.org/10.1016/S1388-2457(02)00106-2>

Polich, J. (1997). EEG and ERP assessment of normal aging. Electroencephalography and Clinical Neurophysiology/Evoked Potentials Section, 104(3): 244-256. <https://doi.org/10.1016/S0168-5597(97)96139-6>

Yordanovaa, J. Y., Koleva, V. N., & Bas ̧ar, E. (1998). EEG theta and frontal alpha oscillations during auditory processing change with aging. *Electroencephalography and Clinical Neurophysiology, 108*: 497-505. <https://doi.org/10.1016/S0168-5597(98)00028-8>
